# 2D, or not 2D? Investigating Vertical Signal Integrity of Tissue Slices

**DOI:** 10.1101/2025.01.13.632601

**Authors:** Sebastian Tiesmeyer, Niklas Müller-Bötticher, Alexander Malt, Leyao Ma, Sergio Marco-Salas, Paul Kiessling, Paul Horn, Adrien Guillot, Louis B Kuemmerle, Frank Tacke, Fabian Theis, Christoph Kuppe, Mats Nillson, Roland Eils, Brian Long, Naveed Ishaque

**Author notes:** these authors contributed equally to the work (Sebastian Tiesmeyer, Niklas Müller-Bötticher). these authors jointly supervised the work (Brian Long, Naveed Ishaque).

## Abstract

Imaging-based spatially resolved transcriptomics can localize transcripts within tissue sections in 3D. Cell segmentation assigns transcripts to cells and precedes annotation of cell function. However, cell segmentation is usually performed in 2D, thus unable to deal with spatial doublets arising from overlapping cells, resulting in segmented cells containing transcripts originating from multiple cell types. Here we present a computational tool called *ovrlpy* that identifies overlapping cells, tissue folds, and inaccurate cell-segmentation by analyzing transcript localization in 3D.

## Main

Spatially resolved transcriptomics (SRT) has revolutionized the study of cellular architecture within tissues^1^. However, despite profiling 5–15 µm thick tissue sections, most available SRT studies ignore the vertical dimension and perform planar projection to interpret their data in a completely flat 2D coordinate space^2–4^. This common practice overlooks the vertical complexity inherent in the tissue, potentially conflating signals from overlapping cells and generating multi-cell ‘doublet’ artifacts^5^. These vertical spatial doublets may result from biological cell associations or technical artifacts, such as tissue folding, posing challenges in accurate cell-typing and downstream analysis^6,7^. While sequencing-based SRT methods measure gene expression in 2D, imaging-based SRT methods can resolve transcript locations in 3D depending on the image acquisition device^8^.

To illustrate the potential impact of projecting 3D spatial tissue data onto a 2D plane, we developed a simplified geometric model (**Supplementary Figure 1**). By simulating spherical cells within dense tissue sections of various thicknesses, we calculated the “signal-to-total” ratio, representing the proportion of signal originating from the target cell versus vertically adjacent structures. For a 10 µm diameter cell in a 10 µm slice section, the model predicts an average signal-to-total ratio of 0.4. This relatively low ratio questions the practice of treating imaging-based SRT data as 2D, instead suggesting that SRT signal will routinely be contaminated by its vertical environment. It is crucial to note that while the quantitative results from this model may not accurately represent all cell types and tissues, the underlying principle of signal contamination in 2D projections likely holds true across diverse cellular morphologies.

To explore the 3D-to-2D projection effect in tissue sections, we developed a strategy to virtually split tissue slices *in silico* into a top and bottom part. Our initial exploration of 3D transcript localizations revealed that measured transcript z position in some data can vary across the sample (**Supplementary Figure 2a**). In this case, a global drift of vertical position along the sample prevents splitting the sample using a naïve global threshold. So, we developed an analysis approach demonstrated on a 10 µm thick mouse brain coronal section dataset profiled on the Xenium platform. First, the transcripts are binned across the x-y coordinates (usually 1 µm bin size) and the mean transcript z-coordinate is calculated per bin. The z-mean is then locally smoothed using a graph-based message passing strategy by repeatedly averaging across neighboring bins. The resulting z-coordinate is used as the splitting point per bin (**Figure 1a**). Using our locally smoothed mean approach to split the sample into virtual subslices results in more comparable top and bottoms sections (**Figure 1b**).

**Figure 1:**
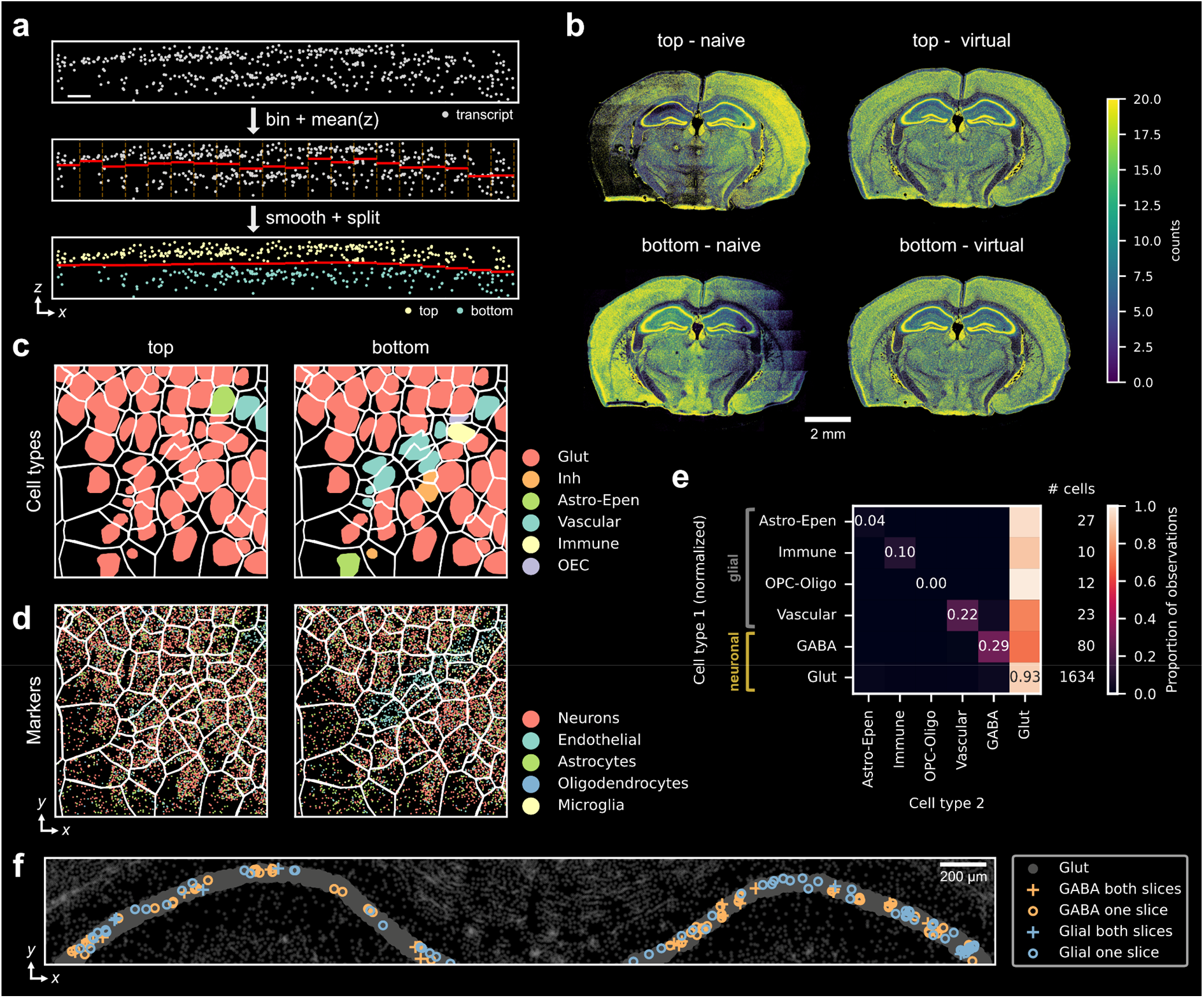
Investigating vertical consistency in tissue sections. **a)** Schematic of the virtual subslicing workflow (simplified for x-axis but operates in 2D i.e. x-y bins). Scale bar corresponds to 1 µm. **b)** Splitting the sample in two using the global mean z-coordinate (naïve) leads to uneven sections compared to the locally smoothed mean (virtual). Counts per 5 µm bin. **c)** MapMyCells annotation of cell segments in the top and bottom virtual sub-slices in the CA1 region. Nuclei are colored and delineated by cell borders. **d)** Transcripts colored by cell-type classes. **e)** Combinatorial matrix of top-bottom observation pairs in virtual hippocampus sub-slices. **f)** Consistency of ‘GABA’ and ‘glial’ assignment of segments across the vertical split.

**Figure 2:**
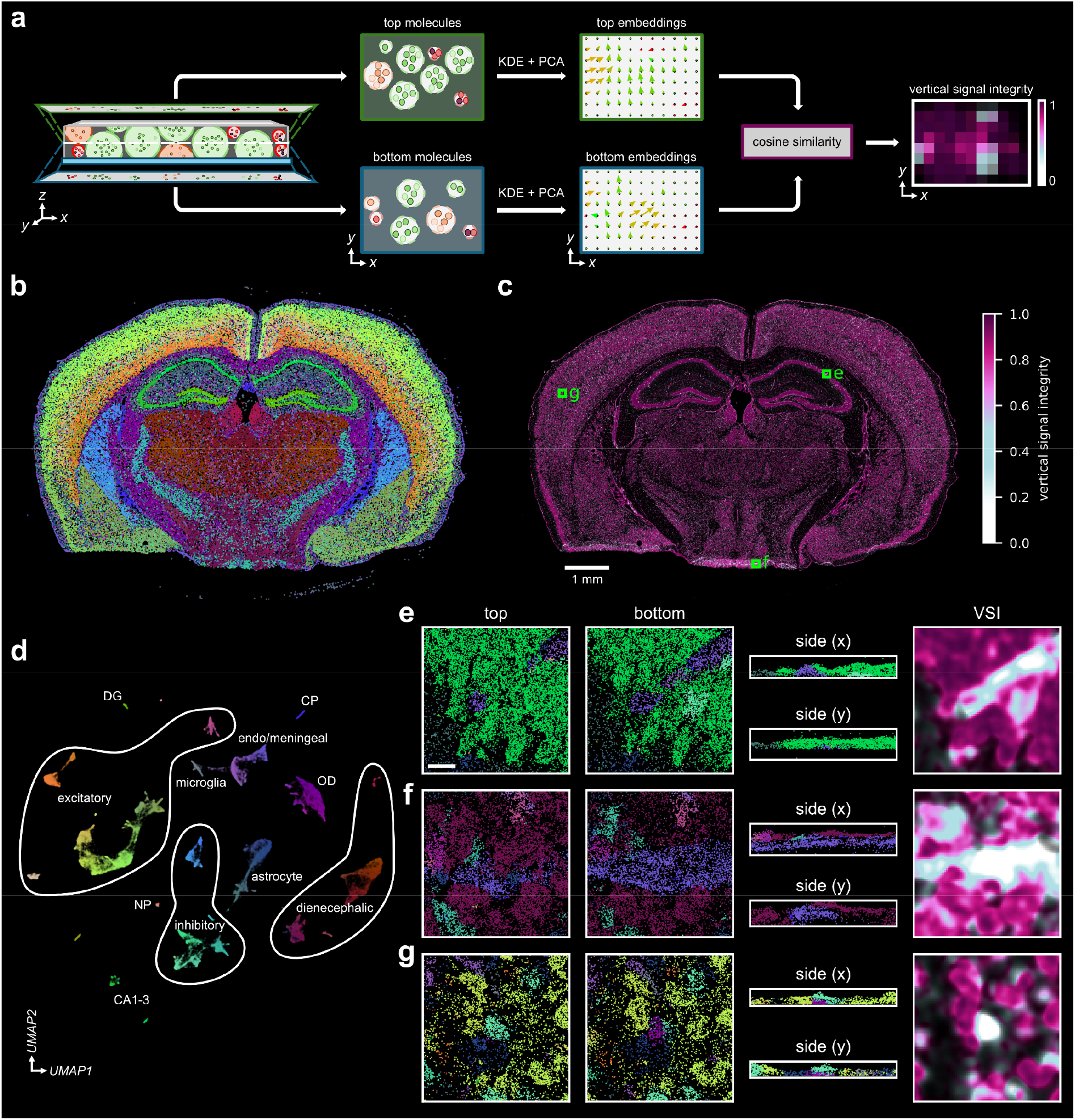
Ovrlpy identifies vertically overlapping cells in imaging-based spatially resolved transcriptomics data. **a)** The ovrlpy vertical signal integrity pipeline. **b)** Color-embedded transcriptome signal at local maxima sampling locations. **c)** Vertical signal integrity (VSI) map. **d)** Segmentation-free, unsupervised transcriptome signal embedding of local maxima. **e-g)** Transcripts colored by the RGB embedding (panel d) and VSI of doublet locations; **e)** for the vessel overlap in the CA1 region highlighted in Figure 1c, **f)** the tissue fold, and **g)** the neocortex. Scale bar corresponds to 15 μm. CP: choroid plexus, DG: dentate gyrus, NP: neural progenitor, OD: oligodendrocyte

Based on our virtual subslicing strategy, transcripts were divided into ‘top’ and ‘bottom’ slices and aggregated according to the original segmentation. The cells in the top and bottom subslices were independently annotated using MapMyCells^9^ (**Figure 1c, d**). Across 162,033 segmented cells, only 78% of cells were consistently assigned the same cell type between top and bottom slices. This was in line with the other sample replicates (80% and 79%). Notably, the top-bottom mismatches implicated that glial cells like astrocytes, oligodendrocytes and vascular cells are especially prone to overlapping, thus leading to vertical spatial doublets in the z-axis (**Supplementary Figure 3**), consistent with the ubiquitous presence of glial cells in central nervous system^10^. We further investigated the hippocampal CA1 region, which is a homogenous, well-structured tissue section consisting mostly of a dense matrix of glutamatergic excitatory neurons interspersed with sparse GABAergic inhibitory neurons and glial cells^11^. Among 152 detected inhibitory/glial cells, only ~20% showed consistent cell-type assignment between top and bottom subslices, while the majority formed vertical doublets with neighboring glutamatergic neurons (**Figure 1e-f, Supplementary Figure 3b**).

To investigate whether the effect is limited to the Xenium technology, we investigated a mouse brain coronal section profiled on the MERSCOPE platform. The global z-drift was less pronounced, with transcripts limited to a few discrete z-coordinates (**Supplementary Figure 2b, c**). Similar to the Xenium sample we observed a vertical assignment consistency of 83% over the entire tissue, and 51% for non-neuronal cells in the hippocampal CA1 region (**Supplementary Figure 4-5**).These results likely overestimate the impact of vertical doublets on cell type mapping in the original data (because the partial cells contain fewer transcripts reducing the cell-typing quality), but the mapping results indicate the presence of vertical contamination of transcriptomic profiles in a cell-type and tissue-dependent manner.

To address vertical spatial doublets in imaging-based SRT data generated from tissue sections, we implemented *ovrlpy* (https://github.com/HiDiHlabs/ovrl.py). Ovrlpy combines the vertical subslicing strategy with an unsupervised and segmentation-free SRT analysis algorithm^12,13^. Ovrlpy generates tissue maps of vertical signal integrity (VSI) by calculating local gene expression similarity between top and bottom virtual subslices (**Figure 2a**). By simulating tissue overlaps *in silico*, we demonstrate that low VSI corresponds to cell overlaps (**Supplementary Figure 6**). Importantly, the ovrlpy algorithm does not rely on or incorporate the simplified geometric model in any way. Our tool’s functionality is independent of cell shape assumptions, making it applicable to tissues with diverse cellular morphologies. Ovrlpy can identify and visualize regions of interest (ROI) for further inspection. The transcriptome signal is embedded both in 2D-UMAP space and in an RGB-color space. Subsequently, the 2D view of transcripts in the ROI is colored by the RGB embedding.

An example application to the Xenium mouse brain data set revealed different types of low-VSI artifacts (**Figure 2b-g, Supplementary Figure 7**). The unsupervised color assignment of transcripts corresponded well to different cell-types. Low-VSI peaks appear throughout the tissue, indicating widespread cell overlaps. The persistently low VSI observed in the meninges, choroid plexus, and capillaries, suggests that cellular morphology and tissue structure play a significant role in determining VSI in these regions. Many of these observations were consistent with the previous segmentation-based analysis (**Figure 2e**). The ventral brain region on the bottom contains several prominent structures, which had low VSI and visual consistency with sample preparation tissue folds (**Supplementary Figure 7**). Furthermore, ovrlpy confirmed vertical doublets involving glial and inhibitory cells in the CA1 region (**Supplementary Figure 8**).

These observations were also replicated when applying ovrlpy to the MERSCOPE mouse brain dataset (**Supplementary Figure 9**), and vertical doublets could be validated in z-stack DAPI images (**Supplementary Figure 10**). Furthermore, we demonstrated that a virtual subslice has an overall higher VSI than the corresponding full sample (**Supplementary Figure 11**) thus validating the intuition that thinner slices should contain less cell overlaps.

To investigate the generalizability of ovrlpy across tissues, we analyzed a publicly available mouse liver dataset profiled on the MERSCOPE platform and were able to generate unsupervised gene expression embeddings consistent with the liver cellular composition^8^. Patterning of the VSI map as well as instance visualizations suggest an increased tendency for liver endothelial cells and stromal cells to form vertical doublets with their immediate environment (typically hepatocytes as well as, in the case of endothelial cells, other building blocks of multilayered vascular structures, **Supplementary Figure 12, 13; Supplementary Results**).

3D segmentation methods offer a potential solution to address vertical spatial doublets for imaging-based SRT data but are limited by technical challenges. Reduced microscopic resolution along the z-axis impedes precise boundary delineation, and cells partially outside the tissue slice pose difficulties for standard model priors as in the case of missing DAPI signal or inapt cell size constraints. To demonstrate ovrlpy’s utility as a cell-segmentation quality control tool, we investigated two 3D segmentation strategies. First, we performed 3D cell segmentation based on transcript locations of the Xenium mouse brain dataset using Baysor^14^ and Proseg^15^. For the obtained segments, we observed a significant correlation of MapMyCells’s reported cell typing correlation coefficient and ovrlpy’s reported VSI score (**Supplementary Figure 14**). However, both Baysor and Proseg struggled to model the tissue fold artifact in the ventral region, failing to correctly segment a number of leptomeningeal cells along a stretch of the folded meninges in regions where the leptomeningeal markers were clearly present. We then investigated a publicly available 3D DAPI segmentation dataset of the mouse brain from Vizgen (**Supplementary Figure 9)**. Less than 7% of the 810 high confidence overlap regions (VSI < 0.5) identified by ovrlpy had overlapping nuclei-segments at the detected location, suggesting limitations in existing 3D nuclear segmentation models’ ability to identify vertically overlapping cells effectively. These findings underscore ovrlpy’s sensitivity to spatial artifacts that may be missed by existing 3D cell segmentation methods and emphasize the urgent need for robust quality control strategies in SRT analysis.

We further investigated the effect of ovrlpy-defined vertical doublets on downstream cell typing analysis. Mean VSI scores were determined across each cell segment, and segments with a mean VSI below 0.7 were marked as vertical doublets. We then annotated all cell segments and inspected their gene expression UMAP embeddings. We found that the doublets were embedded in between the more distinct singlets, as observed with single-cell RNAseq doublet gene expression embeddings^16^. Removing the vertical doublets resulted in an improved signal-to-noise ratio, demonstrated by a clearer separation of cell-types in both the Xenium and MERSCOPE datasets (**Supplementary Figure 15**). Furthermore, we show that ovrlpy can help identify spurious spatial cell-typing that most likely results from gene expression contamination through overlapping cells rather than true cell subtype differences (**Supplementary Figure 16**).

To ensure accessability and usability of ovrlpy, we comprehensively profiled its computational requirements (**Supplementary Figure 17**) and made the package available via PyPI and bioconda.

In conclusion, our analysis offers a practical solution to identify and reduce the artifacts introduced by assuming SRT data to be 2D. Researchers should consider the frequent phenomenon of overlapping structures, vertical spatial doublets and partly-out-of-slice cell fragments when planning experiments, interpreting results, or creating computational analysis tools. The VSI score reflects signal integrity between virtual tissue subslices and can be influenced by *(i)* vertical cell overlap, where transcripts from multiple z-plane cells are captured together, and *(ii)* tissue processing artifacts, such as folds. Furthermore, factors leading to non-uniform distribution of transcripts within individual cells, such as subcellular mRNA localization patterns or uneven tissue sectioning, potentially affect VSI. Therefore, users should examine the cellular composition of low VSI regions to distinguish between these different sources of signal heterogeneity. Furthermore, a number of parameters may influence VSI distributions, making context-specific evaluation of VSI thresholds essential for biological interpretation. While we present multiple lines of evidence that ovrlpy can identify cell overlaps in spatial transcriptomics data and show improved performance over existing 3D segmentation strategies, a systematic evaluation of its performance in light of emerging 3D and multi-modal cell segmentation is warranted. Limitations in current cell segmentation strategies are highlighted by the emergence of tools and strategies to evaluate and mitigate inaccurate segmentation^15,17,18^. To the best of our knowledge, ovrlpy is the only unsupervised tool to focus on potential spatial doublets in the z-axis, that can also be used in conjunction to add a third dimension to tools that focus on 2D quality assurance of segmentations. Ovrlpy is an unsupervised algorithm, thus making it suitable for both known and unknown tissues and cell types. The identified overlaps are plausible given cell-type specificity of transcripts and known cellular organization within tissues. As SRT continues to evolve and is increasingly applied to attack real-world biological problems, tools like ovrlpy will be essential for ensuring the integrity and interpretability of high-resolution spatial data. Ovrlpy is compatible with standard Python SRT analysis infrastructure, provides tutorials of example applications as well as guidelines for optimal model parameter selection and output interpretation.

## Notes

### Competing Interest Statement

The authors have declared no competing interest.

### Summary of Updates

Revised version. Includes clarifications of geometric model limitations; extended clarity and robustness of the virtual subslicing approach; parameter sensitivity analysis; experimental validation with simulations and 3D DAPI stains; analysis on downstream impact of doublet artefacts.

