## Supplementary material for "2D, or not 2D? Investigating Vertical Signal Integrity of Tissue Slices": online methods

### Online methods for Tiesmeyer et. al. 2D, or not 2D? Investigating Vertical Signal Integrity of Tissue Slices

#### Online methods

Data was analyzed using ovrlpy v1.0.1 in Python v3.12.11 with the following main packages; anndata v0.11.4, matplotlib v3.10.3, numpy v1.26.4, pandas v2.1.4, polars v1.31.0, rasterio v1.4.3, scikit-image v0.25.2, scikit-learn v1.7.0, scipy v1.16.0, seaborn v0.13.2, shapely v2.1.1, and umap-learn v0.5.8.

##### Geometric model

We created two geometric statistical noise models of vertical signal in microscopic tissue slices, which we called ‘signal-to-total’ and ‘clean-to-any’. The models are based on simple, solid geometric bodies, which are assumed to emit a uniform transcriptomic signal throughout their volume. The signal is then observed along the vertical z-axis, and the marginal distribution of volumes along the z-axis is investigated as the local signal after vertical collapse.

###### Expected signal-to-total model

The expected ‘signal-to-total’ $E[\text{signal-to-total}]$ answers the question: during 2D analysis, one observes signal from a given cell type $C$ of known diameter $d$, in a slice of known height $h$. What’s the expected portion of my signal stemming from the source cell of interest itself, instead of extracellular/paracellular structures above or below the cell?

To derive the expected signal-to-total, we first define the parameters:

- cell diameter $d$ and radius $r = \frac{d}{2}$
- slice height $h$
- vertical cell position $z$

Assuming $z$ to be equally distributed (the cell can occur at any vertical position with equal probability), the strategy to derive $E[\text{signal-to-total}]$ is:

1. The expected signal-to-total equals the expected volume of a cell inside the sphere divided by the volume of the vertical outline of the sphere inside the slice:
    a. $E[\text{signal-to-total}] = \frac{E[v_{\text{cell x slice}}]}{E[v_{\text{footprint}}]}$
2. Define a number of basic volume formulas:
    a. The volume of a sphere: $v_{\text{sphere}} = \frac{4}{3}\pi r^3$
    b. The volume of a spherical cap, the part of a sphere above a given horizontal plane at height $h_{\text{cap}}$: $v_{\text{cap}} = \frac{\pi h_{\text{cap}}^2}{3}\left(3r - h_{\text{cap}}\right)$

3. Since the cell is expected to occur at the same probability at any elevation $z$, we can determine the expected volume of the source cell as the intersection between the cell and the slice. This scenario is horizontally mirror symmetric, therefore we assume the slice to be centered at $z = 0$, and define $h' = \frac{h}{2}$ as the extent of the slice above the $z$-origin. We then can investigate only cases of the cell occurring in the top half of the slice and assume the overall expected value to be identical.
   a. $E[v_{\text{cell-in-slice}}] = \int_0^{h'+r} v_{\text{sphere}}\, v_{\text{slice}} \frac{dz}{h'+r}$
4. When $r < h'$ (the cell is smaller than the slice), we can differentiate between the cases where the sphere is entirely or partially inside the slice:
   a. $E[v_{\text{cell-in-slice}}] = \int_0^{h'-r} v_{\text{sphere}} \frac{dz}{h'-r}$
   b. $E[v_{\text{cell-out-of-slice}}] = \int_{h'-r}^{h'+r} v_{\text{cap}} \frac{dz}{2r}$
   c. $E[v_{\text{cell}}] = E[v_{\text{cell-in-slice}}] + E[v_{\text{cell-out-of-slice}}]$
   $$= \int_0^{h'-r} \frac{4}{3}\pi r^3 \frac{dz}{h'+r} + \int_{h'-r}^{h'+r} \frac{\pi h_{\text{cap}}^2}{3}\left(3r - h_{\text{cap}}\right)\frac{dz}{2r} = \frac{4}{3}\pi h r^2$$
   where $h_{\text{cap}} = h' - r + z$
5. The observed signal inside the cell's footprint equals the volume of a cylinder of height $h$ with the base radius equal to the widest part of the sphere-slice-intersection:
   a. $E[v_{\text{footprint}}] = \int_0^{h'} \pi h r^2 \frac{dz}{h'} + \int_0^{r} \pi h(\mathrm{r}^2 - \mathrm{z}^2)\frac{dz}{r} = \frac{2\pi h r^2 \cdot (3h+4r)}{3}$
6. According to (1a, 4c, 5a), we can resolve that: $E[\text{signal-to-total}] = \frac{4r}{4r+3h} = \frac{2d}{2d+3h}$

#### Expected clean-to-any model

The expected clean-to-any signal models the probability of an observed vertical signal being entirely clean – that is, the observed cell covering the entire vertical space of the tissue at the observed point.

Trivially, if $d < h$, vertical signal will always be contaminated by extracellular structures above or below the source cell, so $E[\text{clean-to-any}] = 0$.

For the other cases, we can build a similar model to signal-to-total but only consider signal from cells that penetrate the entire tissue slice.

1. The formula for the volume of the footprint of the signal remains the same:
   $$E[v_{\text{footprint}}] = \frac{2\pi h r^2 \cdot (3h + 4r)}{3}$$
2. We now consider only the cylindric portion of the sphere that covers the entire tissue slice from top to bottom. It should be equal to the volume of a cylinder of height $h$ and a base area corresponding to the intersection circle of the sphere with the slice surface.
   a. The intersection area sphere × slice-surface is equal to:
   $A_{\text{vertical-clean}} = \pi\left(r^2 - \left(r - h_{\text{cap}}\right)^2\right)$
   b. The volume of a cylinder of height $h$ above this area is then:
   $$E[v_{\text{clean-signal}}] = \int_0^{r-h'} A_{\text{vertical-clean}}\, dh_{\text{cap}} = 2\pi r(-h + r)^2 - \frac{2\pi(-h + r)^3}{3}$$
3. The quotient of the clean signal and the footprint is used to compute the signal-to-any ratio:

$$E[\text{clean-to-any}] = \left(2\pi r(-h+r)^2 - \frac{2\pi(-h+r)^3}{3}\right) \cdot \frac{3}{2\pi h r^2 \cdot (3h+4r)}$$
$$= \frac{(h-r)^2(h+2r)}{(3h+2r)r^2}$$

#### *In silico* validation of geometric models

To validate the proposed geometric models, we performed computational simulations of the ball-in-slice scenario and quantified the observed signal-to-total and clean-to-any voxel ratios across a range of slice heights and sphere radii. Specifically, slice heights were parameterized from 1 to 25 pixels in increments of 1, while sphere radii were varied from 1 to 50 pixels in increments of 2. A 3D voxel space with a default height of $h = 0$ was initialized. Within this voxel space, a sphere of radius $r$ was positioned iteratively along the $z$-axis at discrete coordinates spanning $[-r, r]$ in steps of 1 pixel. Voxels occupied by the sphere were assigned a value of 1, representing inclusion within the sphere volume.

The resulting 3D voxel space was collapsed into a 2D matrix by calculating the mean voxel value along the $z$-axis, effectively simulating a vertical projection of the sphere's occupancy. In this 2D matrix, pixels with values greater than 0 were categorized as "any observations", representing regions where the sphere intersected the slice. Pixels with values exactly equal to 1 were categorized as "clean signals", indicating complete occupancy by the sphere. These classifications enabled the computation of the signal-to-total and clean-to-any voxel ratios across the parameter space.

To ensure interpretability in a biological context, sphere diameters, rather than radii, were reported as the primary geometric parameter in alignment with conventions in microanatomical studies. The observed ratios are displayed alongside the theoretical values derived from the proposed statistical models (**Supplementary Figure 1**).

#### *In silico* vertical subslicing

Accurate vertical alignment of imaging-based spatial transcriptomics data is essential to mitigate technical and procedural artifacts, including uneven tissue sectioning, slanted slide positioning, gravitational settling on glass substrates, and distortions from optical guidance systems[1]. To address these challenges, *ovrlpy* calculates the vertical center of mass (COM) of mRNA molecule coordinates within a horizontal grid (typically 1 µm in width). By default, the COM is defined as the mean $z$-elevation of molecules in each grid segment, although users may specify alternative statistical measures, such as the median, to accommodate dataset-specific features.

To minimize abrupt transitions and excessive fluctuations in the COM calculation, *ovrlpy* incorporates an optional smoothing algorithm based on graph-based message-passing. This method computes a more integrated local $z$-elevation by considering the vertical positions of molecules in neighboring grid pixels. For each grid pixel $i$, a neighborhood graph $G$ is defined using the 4-connected neighbors. The initial COM is calculated for each pixel and iteratively updated by averaging the COM values of neighboring pixels:

$$COM_i^{(t+1)} = \frac{1}{|N_i|} \sum_{j \in N_i} \frac{COM_j^{(t)} + COM_i^{(t)}}{2}$$

where $N_i$ represents the neighboring pixels of $i$, and $t$ denotes the iteration. By default, the process runs for 20 iterations, yielding a smoothened COM profile while maintaining alignment fidelity.

After refinement, the tissue z-axis is reoriented to center the COM at the $z$-origin, enabling uniform virtual slicing. This alignment preserves the original (x, y) coordinates of molecules and maintains the relative vertical distances between molecules sharing the same (x, y) position.

### Vertical signal integrity

#### Gene expression model

*Ovrlpy* combines the subslicing routine described above with a segmentation-free, unsupervised signal embedding algorithm inspired by SSAM[2]. This approach determines vertical signal integrity in the tissue across two *in silico* virtual subslices. The algorithm utilizes a Gaussian kernel density estimation (KDE)-based gene expression model to sample gene expression data from SRT datasets at locations with high gene expression signal. Sampling points with high signal intensity, referred to as "pseudo-cells" in the *ovrlpy* interface, are identified using a local maxima detection algorithm applied to a Gaussian KDE of the entire SRT signal:

$$f(x) = \sum_{i=1}^{n} K\left(\frac{x - x_i}{\sigma}\right)$$

where:

- $X = \{x_1, x_2, \dots, x_n\}$ is the set of observed transcript molecules
- $K$ is the Gaussian kernel function
- $\sigma > 0$ is the bandwidth parameter controlling the smoothness of the KDE

Local maxima $M$ across all genes are then detected at locations $x^*$, where the gradient of the kernel density is 0, the curvature (as per the Hessian matrix of second derivatives $H$) is definite negative, and the value of the kernel density exceeds a threshold parameter $\tau$:

$$\mathrm{M} = \left\{ x^* \in \mathbb{R}^d \mid f(x^*) > \tau \wedge \nabla f(x^*) = 0 \wedge H\big(f(x^*)\big) < 0 \right\}$$

A filter is applied to the local maxima, where all maxima occurring within a minimum distance $d_{min}$ of a larger local maximum are discarded, making sure that the identified pseudo-cells are at least $d_{min}$ apart.

The Gaussian kernel bandwidth $\sigma$**,** the minimum distance between local maxima $d_{min}$ and the KDE sampling threshold $\tau$ are externally defined parameters. The minimum signal threshold $\tau$ ensures that sample locations with low signal strength (regarded as extra-cellular) are removed, making sure that isolated molecules are excluded as pseudo-cell locations.

After filtering, gene expression profiles are generated by applying KDE to transcript locations of individual genes and sampling at the determined local maxima to create a pseudo-cell-by-gene count matrix $C \in \mathbb{R}^{m \times p}$:

$$C_{j,g} = \sum_{x \in X_g} K\left(\frac{x_j^* - x}{\sigma}\right)$$

where $G = \{g_1, g_2, \dots, g_p\}$ is the set of genes, $X_g = \{x_1, x_2, \dots, x_n\}$ are the spatial transcript locations for gene $g$, and $C_{j,g}$ a gene-pseudo cell expression matrix.

Principal component analysis (PCA) is applied to the gene expression matrix to capture patterns of recurring gene expression profiles in a lower dimensional latent space $Z \in \mathbb{R}^{m \times k}$:

$$Z = (C - \mu)V$$

where $\mu$ is the mean vector of $C$, $V = [v_1, v_2, \dots, v_k]$ denotes the top $k$ eigenvectors of the covariance matrix of $C$. The number of PCA components $k$ can be adjusted by the user.

#### Vertical signal integrity maps

The gene expression model described above is used to calculate vertical signal integrity (VSI) across the sample. To compute VSI maps, transcripts in the tissue are divided into top and bottom slices using the mean center of mass (COM) vertical subslicing strategy. For the top and bottom *in silico* slices, a gene expression vector field is generated using 2D KDE with the same kernel bandwidth parameter as in the cell-gene sampling algorithm:

$$C_{\mathrm{top},g}(l) = \sum_{x \in X_{\mathrm{top},g}} K\left(\frac{l - \mathrm{x}}{\sigma}\right)$$

$$C_{\mathrm{bottom},g}(l) = \sum_{x \in X_{\mathrm{bottom},g}} K\left(\frac{l - \mathrm{x}}{\sigma}\right)$$

where $l$ is the location in space at which the kernel is computed.

The resulting gene expression vector fields for the top and bottom slices are projected into PCA space using the principal component vectors $V$ determined previously, resulting in a top and bottom vector field in reduced, "pseudo-cell type" space:

$$Z_{\mathrm{top}}(l) = \left(C_{\mathrm{top}} - \mu_{\mathrm{top}}\right) V$$

$$Z_{\mathrm{bottom}}(l) = \left(C_{\mathrm{bottom}} - \mu_{\mathrm{bottom}}\right) V$$

*Ovrlpy* then calculates a VSI map by measuring the cosine similarity between corresponding top and bottom vectors:

$$\mathrm{VSI}(l) = \cos\theta = \frac{Z_{\mathrm{top}}(l) \cdot Z_{\mathrm{bottom}}(l)}{\left\|Z_{\mathrm{top}}(l)\right\| * \left\|Z_{\mathrm{bottom}}(l)\right\|}$$

where

- $Z_{\mathrm{top}}(l) \cdot Z_{\mathrm{bottom}}(l)$ is the dot product between the two expression vectors
- $\|Z\|$ is the Euclidean norm of $Z$

#### Unsupervised gene-expression embeddings and color assignment

To contextualize the VSI analysis, *ovrlpy* provides functionality to visualize three-dimensional color-embedded transcriptome signals. The 3D-local maxima gene expression samples $C$ generated during VSI map creation are used to construct two non-linear gene expression UMAP embeddings. The first embedding is a "traditional" two-dimensional UMAP while the second is a three-dimensional RGB space embedding to map color values to recurring gene expression patterns (i.e. cell types).

#### 2D UMAP embedding

For the 2D UMAP embedding, PCA-transformed local maxima samples obtained during the VSI map generation $Z$ are reused. A 2D UMAP manifold is created for $Z$ at a minimal embedded distance ('min_dist') of 0 and a neighborhood graph of size 20 ('n_neighbors') by default.

#### 3D UMAP RGB color embedding

The 3D UMAP RGB embedding follows the same strategy as the 2D UMAP embedding but includes additional steps to ensure that the RGB color space is utilized effectively.

A 3D UMAP embedding is generated from the PCA-transformed gene expression data $Z$, with each UMAP dimension corresponding to one color channel in the RGB space. The UMAP algorithm is configured with a minimum distance parameter of 0 and a neighborhood size of 10 by default to capture fine-grained relationships. Before embedding, the vectors in $Z$ are L2-normalized to standardize their magnitudes.

Since UMAP embeddings are unconstrained in range, and RGB values must fall within the $[0, 1]$ interval, a post-processing step ensures proper scaling and alignment of the 3D UMAP outputs:

1. PCA is applied to the 3D UMAP embedding to identify the axes of largest variance in the resulting RGB space.
2. A linear transformation is applied to rotate the embedding so that the principal axes align with the RGB diagonals and scale the values to fit within the $[0, 1]$ range. This ensures efficient and interpretable use of the RGB space.

#### Visualizing a region of interest

The microanatomy of individual sample parts can be visualized by displaying individual transcripts in space, colored according to each transcript's RGB-embedded gene expression vectors. Similar to local maxima sample extraction, the immediate gene expression environment of the transcripts is aggregated using a 3D KDE (see 'Vertical signal integrity'). Contrary to local maxima sampling, KDE values are now sampled at all transcript locations. The expression vector for each transcript is dimensionality reduced using the pre-fitted PCA model as described above, L2-normalized, and projected into the RGB space using the pre-fitted UMAP model. The obtained values are subjected to the same linear transformations as the initial UMAP RGB data, and the output is clipped between $[0, 1]$ to create valid RGB values which are used to color each transcript.

### Identification of vertical doublets as local minima of vertical signal integrity

To facilitate the interpretation of incoherence in tissue samples, *ovrlpy* includes a function to identify individual overlap events by detecting local minima in the VSI map. This functionality enables the identification of regions where vertical gene expression profiles exhibit the lowest coherence, indicative of potential vertical doublets or overlapping structures.

The local minima detection algorithm is an inverted local maxima detection, identifying minima within a user-specified radius on the VSI map. To ensure meaningful results, the function incorporates a threshold parameter to filter out regions with low gene expression vector field norms, which may lead to unreliable and noisy predictions. The identified local minima of vertical signal integrity are prime candidates for further visualization and analysis. These regions allow users to generate targeted visualizations, highlighting potential segmentation problems or areas of overlapping cell types.

##### Evaluation of vertical doublet filtering

To evaluate the impact of applying *ovrlpy's* VSI scores as a filter for segmentation-based cell typing, we conducted a downstream analysis using dimensionality reduction techniques. Gene expression signatures within cell segments or nuclear segments were first subjected to PCA, retaining the first 30 components for further processing. Two-dimensional UMAP embeddings were then generated from these components to visualize the transcriptomic landscape. Cell types were assigned using the MapMyCells[3] online portal with the "10x Whole Mouse Brain (CCN20230722)" reference dataset using the correlation mapping algorithm.

To apply the *ovrlpy* vertical doublet filter, VSI scores were aggregated across each nuclear segment by calculating the mean VSI of all pixels contained within the segment. Segments with an averaged VSI score below 0.7 were excluded, effectively filtering out low-integrity regions likely to contain vertical spatial doublets. PCA and UMAP analyses were subsequently repeated on the filtered dataset, using identical parameter settings to ensure consistency.

#### 10x Genomics Xenium mouse brain

To demonstrate *ovrlpy*, we investigated the publicly available Xenium mouse brain dataset[4]. This dataset includes 62,384,369 detected transcript molecules across 248 selected cell marker genes. The tissue sample was prepared using fresh-frozen mouse brain sections approximately 10 µm thick and processed according to Xenium's 2023 standard analysis protocol. The protocol involves a 2D DAPI-based cell segmentation approach, identifying 162,033 segmented cells.

The *ovrlpy* pipeline was applied to the Xenium dataset using a KDE bandwidth of 2.5 µm (default), 30 principal components (default), and the min_transcripts in the analyse function set to 20 to reduce noise outside the tissue.

##### Comparative analysis of virtual top-bottom slices using supervised annotation

To compare the concordance of the top and bottom split we investigated cell-type annotation generated by MapMyCells. All nuclear transcripts were split into a top and bottom slice based on the virtual subslicing strategy and aggregated according to the original Xenium segmentation. The cell-by-gene counts were exported as AnnData object and submitted to the MapMyCells[3] online interface for cell-type annotation using the "10x Whole Mouse Brain (CCN20230722)" reference dataset and the correlation mapping algorithm. From the predicted cell types of both virtual slices, a combinatorial cell-type matrix was constructed to capture all observed combinations of assigned cell types for the same cell in the top and bottom counterparts. This matrix was normalized row-wise to account for variability in the number of cells across cell types.

##### Vertical signal integrity in the CA1 region

A rectangular window encompassing the CA1 hippocampal region was manually selected based on its characteristic dense architecture. To create a smooth spatial mask of the CA1 region, spatial signal integration was performed using a Gaussian KDE with a bandwidth of 15 µm applied to all transcript molecules followed by density-based filtering (threshold of 2.5) to retain only the dense regions of the CA1 segment.

Subsequently, the comparative analysis of virtual top-bottom slices described above was repeated, restricting the analysis to cells with centroids within the CA1 segment. The VSI score was sampled at the centroid of each cell segment.

#### Artificial folding artifact

An artificial folding artifact was created by stacking two histologically diverse regions *in silico*, simulating a tissue fold. Two square regions (edge length 2 mm) were manually selected from different tissue domains (cortex and thalamus), therefore assumed to be transcriptionally diverse. A mock fold-over was produced by placing the two cut-outs on top of each other and generating a vertical signal integrity map for the stacked tissue, using the same parameterization as for the single tissue slice.

#### 3D cell segmentation of sample preparation artifacts in Xenium mouse brain data

We tested 3D transcript segmentation algorithms (Baysor and Proseg) for their ability to resolve vertical spatial doublets within the prominent fold-over artifacts in the anterior region of the Xenium sample. The resulting gene-cell count matrix was annotated using the MapMyCells online tool with the "10x Whole Mouse Brain (CCN20230722)" reference dataset using the correlation mapping algorithm.

The meningeal markers *Aldh1a2*[5,6], *Col1a1*[7], *Fmod*[8], and *Slc13a4*[9] present in the data set were used to identify vascular and leptomeningeal cell (VLMC) segments. MapMyCells' annotations were compared to segments to evaluate the detection of VLMC annotations within folded tissue regions by manually assigning cells to folded and unfolded regions. For validation, vertical signal integrity values computed by *ovrlpy* at segmentation-identified cell centroids were compared with correlation coefficients (of the "subclass") from MapMyCells' annotations and the Pearson correlation coefficient and p-value for the two metrics calculated using the scipy.stats.pearsonr function using 10,000 permutations.

##### *Baysor*

3D segmentation with Baysor[10] (v0.7.1) was generated by employing Xenium-provided DAPI-based nuclei segmentation as a prior, set at a segmentation confidence threshold of 0.5. Other parameters included a minimum of 10 molecules per gene, 50 molecules per cell, and a scale factor of 5.

##### *Proseg*

To generate a 3D segmentation using Proseg[11] (v2.0.4), the preset Xenium profile was used (--xenium) and the excluded genes were set to remove all features with 'BLANK' or 'NegControl' prefixes.

#### Vizgen MERSCOPE mouse brain

The Vizgen MERSCOPE mouse coronal brain dataset[12] (Slice 2, Replicate 1) consists of a thin FFPE-prepared section. This dataset includes 3D cell segmentation derived from nuclear staining and consists of 48,574,461 detected transcripts, assigned to 438 marker genes. To obtain the assigned cell-ID per transcript the assignment of transcripts was rerun using the original segmentation. Briefly, using the Vizgen post-processing tool[13] (vpt v1.3.0) the segmentation masks were converted to a suitable format (convert-geometry), transcripts assigned to cells (partition-transcripts), and cell metadata generated (derive-entity-metadata).

The *ovrlpy* pipeline was applied to the Vizgen MERSCOPE dataset with all key parameters including the KDE bandwidth, number of principal components (PCs), and min_transcripts at their default.

#### Comparative analysis of virtual top-bottom slices using supervised annotation

Comparison of a virtual top-bottom split was performed analogous to the Xenium brain sample using all transcripts instead of only nuclear assigned transcripts as this information was not available for the MERSCOPE dataset.

#### Verifying vertical signal integrity in CA1

The analysis of the CA1 region was performed as described for the Xenium dataset.

#### Artificial folding artifact

An artificial tissue folding artifact was generated as described for the Xenium mouse brain sample but using 500 µm sized regions. Given the tearing artifacts present in many regions of the sample, areas with sufficiently intact tissue density were prioritized for selection. The selected regions stemmed from the cortex and a region covering the lateral brain including parts of the amygdala, the globus pallidus, and the internal capsule.

#### Vizgen MERSCOPE 3D segmentation of *ovrlpy* vertical doublets

To evaluate the performance of the imaging-based 3D segmentation provided with the MERSCOPE dataset, the segments for all z-planes were visualized alongside the RGB-embedded transcriptome and the VSI map for the corresponding *ovrlpy*-identified vertical doublet.

### Vizgen MERSCOPE mouse liver

The MERSCOPE liver showcase dataset[14] analyzed in this study is a mouse liver dataset (Liver1, Slice1), targeting a gene panel of 347 genes. For the analysis, the dataset was cropped to exclude sample edges and obtain a region measuring 7 mm × 7 mm with 276,188,705 transcripts.

The *ovrlpy* algorithm was applied to the MERSCOPE liver dataset with parameters optimized to capture the thinner, multilayered vascular structures of liver tissue. The KDE bandwidth was set to 1.7 µm, the number of PCs to 15, and the minimum distance between two local maxima to 6 µm.

#### Modified RGB embedding strategy

To investigate cellular heterogeneity in non-parenchymal regions, a second RGB embedding was generated specifically for these regions. Leveraging the separation of parenchymal hepatocytes from non-parenchymal cells in the 2D UMAP embedding, local maxima located in the non-parenchymal regions were extracted and processed independently through the standard *ovrlpy* UMAP and RGB embedding pipeline.

The resulting UMAP and RGB embeddings yielded more distinct clusters of rare cell types. These clusters were annotated using marker gene expression profiles, referencing the Liver Cell Atlas[15], and validated through anatomical expression patterns with expert input from liver biologists. For the remaining analysis, we employed an RGB rendering strategy of first separating expression vectors based on their clustering position on the 2D UMAP into a ‘parenchymal’ and ‘non-parenchymal’ category. Parenchymal expression vectors were rendered in grayscale using their RGB intensity, and non-parenchymal cells were subsequently subjected to the second, non-parenchymal RGB embedding model and displayed using the resulting RGB colors.

##### Artificial folding artifact

An artificial tissue folding artifact was generated as described for the Xenium mouse brain sample. The liver sample offered less opportunity for selecting two regions with different expression profiles due to hepatocyte heterogeneity dominating the UMAP embedding. Therefore, two smaller square regions (edge length 800 µm) were selected from different tissue domains; from the portal triad and from the parenchyme.

#### MERFISH mouse hypothalamus

To evaluate the effect of vertical doublets on spatial cell typing we used the MERFISH mouse hypothalamus from Moffitt *et al.*[16] analyzed by Singhal *et al.*[17] Ovrlpy was run on the MERFISH dataset using 15 principal components (other parameters at the default). To evaluate the VSI of MOD-gm and MOD-wm subtypes (as annotated by Singhal *et al.*) cells not annotated as 'Ambiguous' were selected and VSI scores inside the corresponding segmentation masks with a signal strength above 3 were used.

The scRNAseq reference dataset from Moffitt *et al.* was processed as described by Singhal *et al*. Briefly, only cells with less than 20% mitochondrial counts and more than 1,000 detected genes were kept. The counts were normalized to 10,000 per cell, log-transformed, and z-scaled.

#### Benchmark

To evaluate the computational performance of *ovrlpy* we used the Xenium Prime 5K mouse pup dataset[18]. The genes were downsampled to 1,000 (unless otherwise specified) resulting in 125,756,304 transcripts and an area of 272.6 mm². The runtime and memory usage were evaluated using 8 threads (unless otherwise specified) for varying number of genes (250, 500, 1000, 5010), total area (25%, 50%, 75%, 100%), and number of threads (1, 2, 4, 8, 16). Runtime (ElapsedRaw) and memory usage (MaxRSS) were taken from slurm's job metadata after successful termination.

### Ethical approval

Not applicable.

### Competing interests

The authors declare that they have no competing interests.

### Acknowledgements

The ovrlpy project was conceptualized during the de.NBI BioHackathon SpaceHack project in Lutherstadt-Wittenberg (December 2022), we thank the organizers of and participants in the de.NBI BioHackathon SpaceHack project. We thank Tillmann Rheude for establishing an analogy of 2D analysis of tissue sections to the flat earth conspiracy. This research has received funding from the Federal Ministry of Education and Research of Germany in the framework of SAGE (project number 031L0265) and CNAScope (01KD2443), the German Research Foundation in the framework of CRC/TR 412 (35081457), and the Project grant (2024-02533) from the Swedish Research Council. We would like to thank the GESTALT community for positive engagement and ideas. We would like to thank the following people for useful discussion and sharing data on tissue preparation and other artifacts: Heather C. Etchevers, Aix Marseille University, for the Visium mouse heart samples with folds, smears, and slippage; Thomas Conrad, Berlin Institute of Health at Charité, Francis Baumgartner and Ulrich Keller, Charité – Universitätsmedizin Berlin, for the Xenium human uveal melanoma sample with detachment; Olivier Raineteau, Guillaume Marcy and Cyril Dégletagne, Cancer Research Center of Lyon, for the Xenium mouse brain samples with tissue folds; Marco Grillo and Christoffer Mattsson Langseth, Stockholm University, for the ISS and Xenium multiple sclerosis samples with signal smearing.

### Authors' contributions

NI conceived and designed the study. ST, NMB with some assistance from AM implemented the ovrlpy package. ST, NMB, AM, LM, SM-S, LBK, NI performed data analysis. ST, NMB, BL, NI interpreted the brain sample analyses. ST, PK, PH, AG, CK, NI interpreted the liver sample analysis. ST and NI wrote the manuscript. All authors proofread and corrected the manuscript. All authors contributed to the article and approved the submitted version.

### Code & Data Availability

#### Data availability

This study used publicly available SRT datasets. 10x Genomics Xenium datasets were downloaded from https://www.10xgenomics.com/datasets/fresh-frozen-mouse-brain-replicates-1-standard and https://www.10xgenomics.com/datasets/xenium-prime-ffpe-neonatal-mouse. Vizgen MERSCOPE datasets were downloaded from https://info.vizgen.com/mouse-brain-map and https://info.vizgen.com/mouse-liver-data. The MERFISH hypothalamus dataset was downloaded; metadata from the original publication (https://doi.org/10.1126/science.aau5324), segmented cells from https://doi.org/10.5061/dryad.8t8s248, transcript information and segmentation masks from Zenodo (https://zenodo.org/records/3478502), and the single-cell RNA-seq reference dataset from GEO (GSE113576). The brain reference dataset used for generating signatures was downloaded from the Allen Brain Map (https://portal.brain-map.org/atlases-and-data/rnaseq/mouse-whole-cortex-and-hippocampus-10x). The liver reference signatures were generated from the Mouse StSt dataset of the Liver cell atlas (https://www.livercellatlas.org/). Data generated as part of this study and necessary to reproduce results are available on Zenodo (https://zenodo.org/records/14226546).

#### Software availability

The ovrlpy package is available as free and open-source software on GitHub with a permissive MIT license: https://github.com/HiDiHlabs/ovrl.py. The package can also be downloaded from PyPI (https://pypi.org/project/ovrlpy/) and Bioconda (https://anaconda.org/bioconda/ovrlpy). We provide a repository with Jupyter Notebooks for reproducing all results and figures of this study (https://github.com/HiDiHlabs/ovrlpy-publication).
