## supplementary for "2D, or not 2D? Investigating Vertical Signal Integrity of Tissue Slices"

### Supplementary information for Tiesmeyer et. al.; 2D, or not 2D? Investigating Vertical Signal Integrity of Tissue Slices

### Supplementary Results

#### Verifying 3D UMAP RGB color embeddings

To validate the UMAP color embeddings generated by *ovrlpy*, particularly in regions identified as vertical spatial doublets, we utilized a curated list of cell-type marker genes published alongside the Xenium mouse brain dataset. This list contains 248 marker genes categorized into broad cell types: 'Astrocyte', 'Endothelial', 'Microglia', 'Neuronal', and 'Oligodendrocyte'.

To assess whether *ovrlpy's* RGB embeddings accurately capture biological cell types, RGB-embedded visualizations of tissue microanatomy were compared to the known marker gene information. Regions with overlapping cell types were evaluated by selecting 80 µm windows centered on *ovrlpy*-identified vertical doublets. For each window, transcripts were colored *i)* using *ovrlpy's* RGB embedding, which encodes local gene expression profiles into the RGB color space and *ii)* according to their cell types based on the marker gene categories. This comparison enabled a qualitative confirmation of the correspondence between *ovrlpy's* RGB embeddings and the expected cell type-specific expression patterns (**Supplementary Figure 7**).

#### MERSCOPE mouse liver

The liver tissue's composition, predominantly hepatocytes (~80% of liver mass[1]), posed challenges for automated color embedding. The RGB embedding captured heterogeneity within the hepatocyte population (e.g., periportal versus perivenal zonation) but provided limited representation of rare non-parenchymal cell types, such as endothelial cells, Kupffer cells, immune cells, fibroblasts, and stellate cells (**Supplementary Figures 12, 13**).

The gene expression embeddings confirmed a highly imbalanced cell-type distribution, being predominantly composed of hepatocytes. Moreover, the sample contains structures identified as central veins as well as portal areas, including branches of the portal vein, hepatic artery, and biliary duct. The hepatocyte UMAP embeddings effectively captured the zonation of the liver parenchyma, from periportal to pericentral hepatocytes.

Liver sinusoids are lined with a thin layer of endothelial cells. However, due to their exceptionally thin morphology, these endothelial cells appear to be difficult to retrieve in the transcriptomic domain except for their nuclei, which are sometimes captured on the sinusoid edges. Kupffer cells, the resident macrophages of the liver, line the sinusoids in a similar pattern. Due to their close spatial association with hepatocytes, both cell types have a potential risk of systematic doublet formation.

The *ovrlpy* VSI map reveals a dense and homogeneous liver parenchyma, which is populated by vertical doublets with endothelial and with Kupffer cells. Moreover, the venous structures are salient and show discernible low integrity at nearly all instances. The reason is the laminar build-up of the vascular wall containing endothelial cells, fibroblasts, and smooth muscle cells which, when not cut perfectly perpendicular, form systematic overlapping structures on a large scale, making endothelial cells, stromal cells, and smooth muscle cells especially prone to doublet formation.

### Supplementary Figures

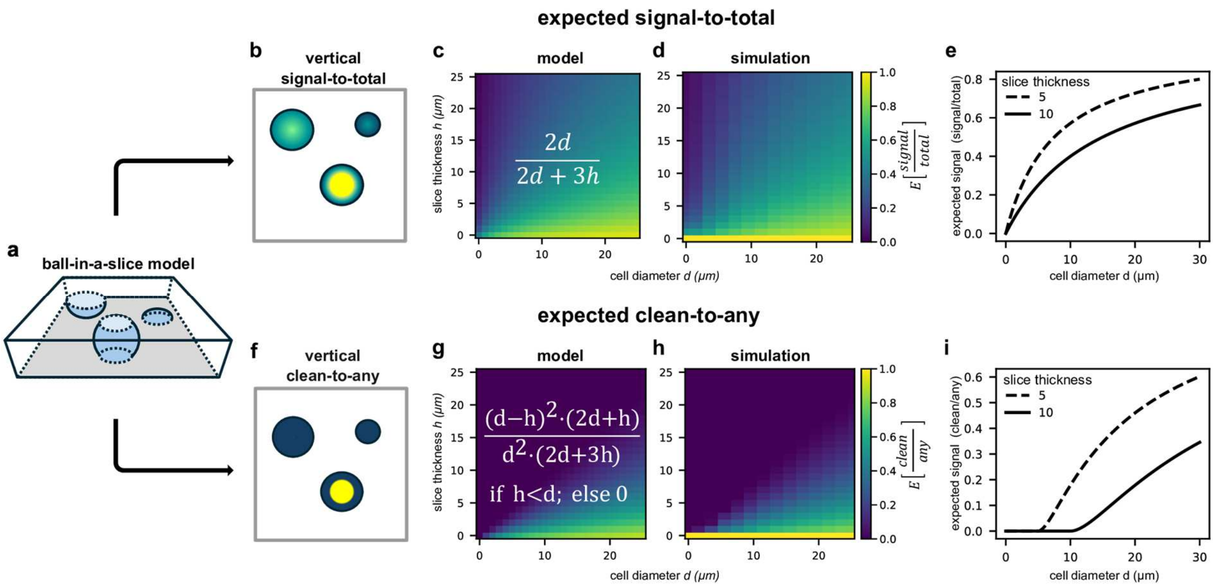

**Supplementary Figure 1: Statistical models of overlapping cells in sliced tissue sections**. **a)** Illustration of a ball-in-slice model, with same diameter balls placed at different elevations in a vertically delimited slice. **b)** Visual interpretation of "signal-to-total" model, as the vertical portion of slice inside the ball. **c)** Modelled expected signal-to-total for different slice heights and cell diameters. **d)** Expected signal-to-total derived by simulation. **e)** Expected signal-to-total model for slice thicknesses of 5 and 10 µm. **f)** Visual interpretation of "clean-to-any" model, as the portion of observed signal without any vertical contamination from outside the ball. **g)** Modelled expected clean-to-any for different slice heights and cell diameters. **h)** Expected clean-to-any derived by simulation. **i)** Expected clean-to-any model for slice thicknesses of 5 and 10 µm.

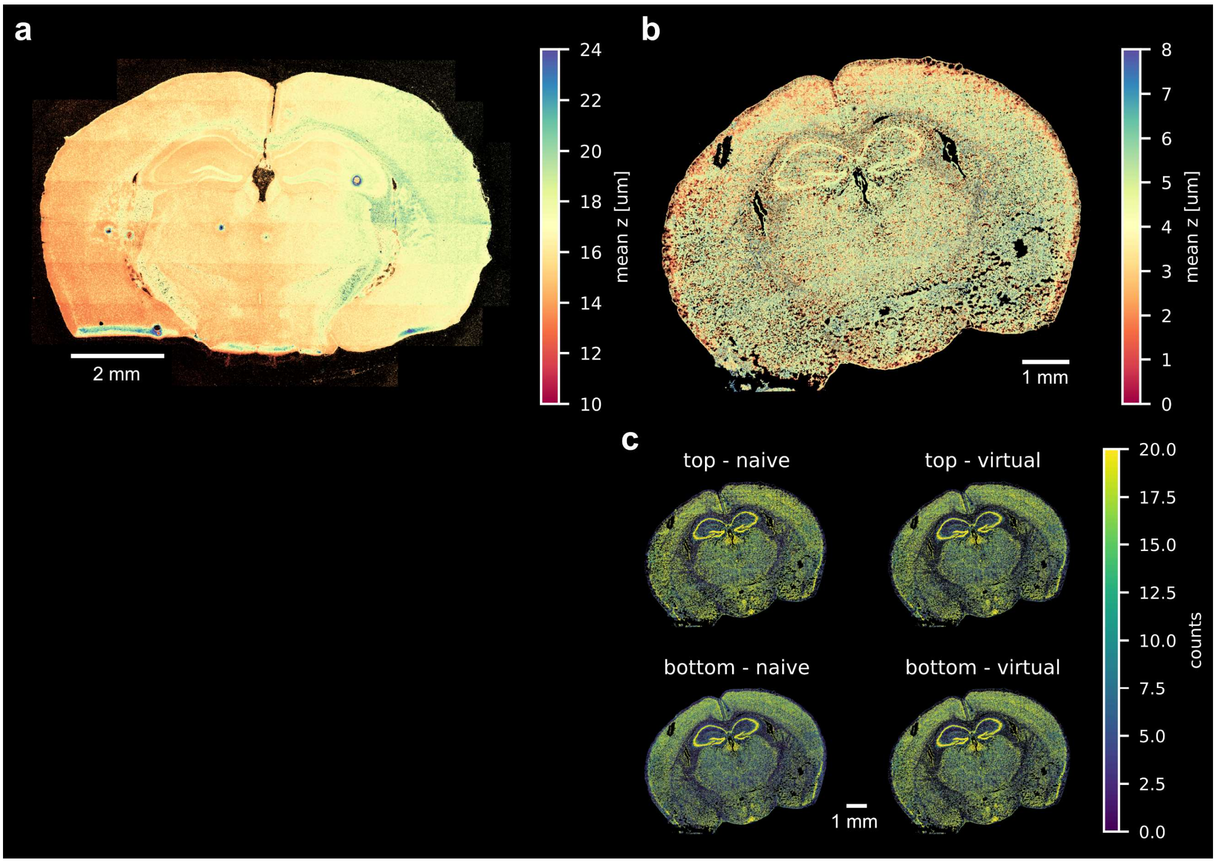

**Supplementary Figure 2: Virtual subslicing overcomes limitations of global z-axis thresholds.** Mean z-coordinate of transcripts (5 µm bins) for **a)** the Xenium mouse brain sample show major global shifts and **b)** minor artifacts for the MERSCOPE mouse brain sample. **c)** Splitting the MERSCOPE sample using the locally smoothed mean z-coordinate (virtual) compared to the global mean (naïve) leads to reduced global artifacts in count density (5 µm bins).

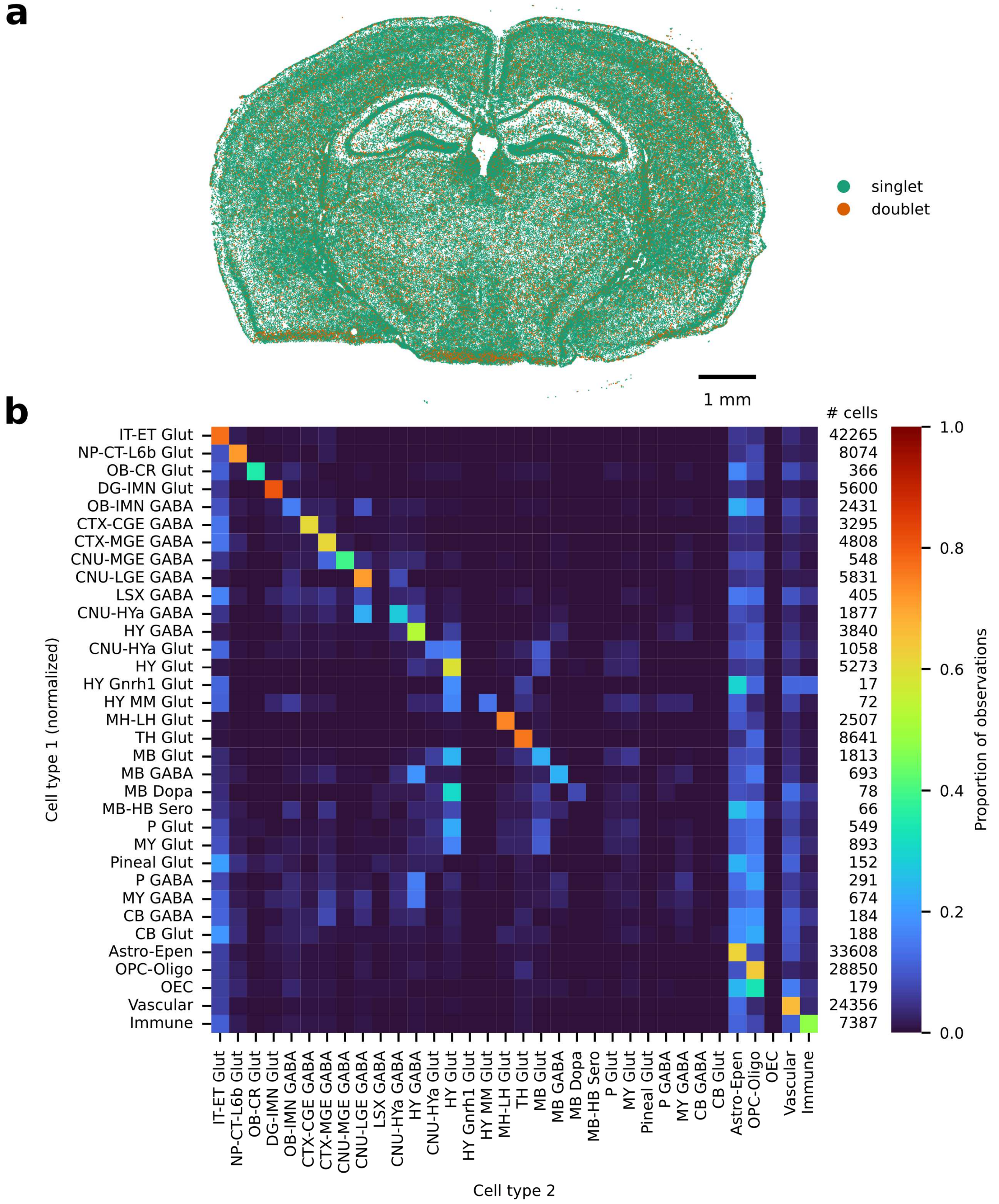

**Supplementary Figure 3: Supervised annotation of virtual subsliced cell segments implicates widespread presence of vertical doublets in the Xenium mouse brain dataset**. **a)** Map of cell segment centroids marked as singlets or vertical doublets depending on their consistency of the MapMyCells assigned class across two virtual vertical subslices. Vertical doublets appear widespread over the tissue, but are more concentrated on the lower, ventral part of the sample. **b)** Cell type-specific observed vertical doublets (row-normalized).

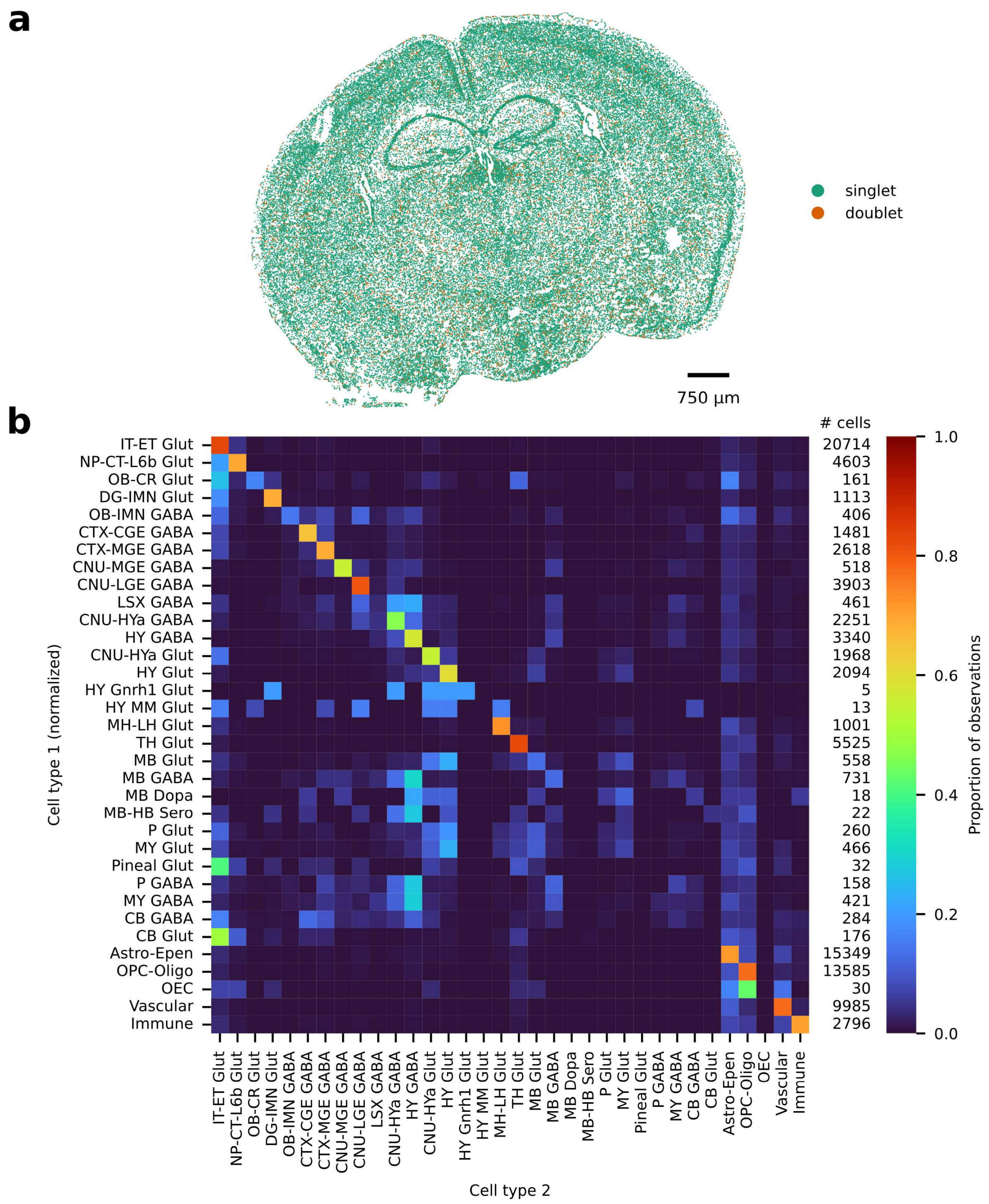

**Supplementary Figure 4: Supervised annotation of virtual subsliced cell segments implicates widespread presence of vertical doublets in the MERSCOPE mouse brain dataset**. **a)** Map of cell segment centroids marked as singlets or vertical doublets depending on their consistency of the MapMyCells assigned class across two virtual vertical subslices. Vertical doublets appear widespread over the tissue. **b)** Cell type-specific observed vertical doublets (row-normalized).

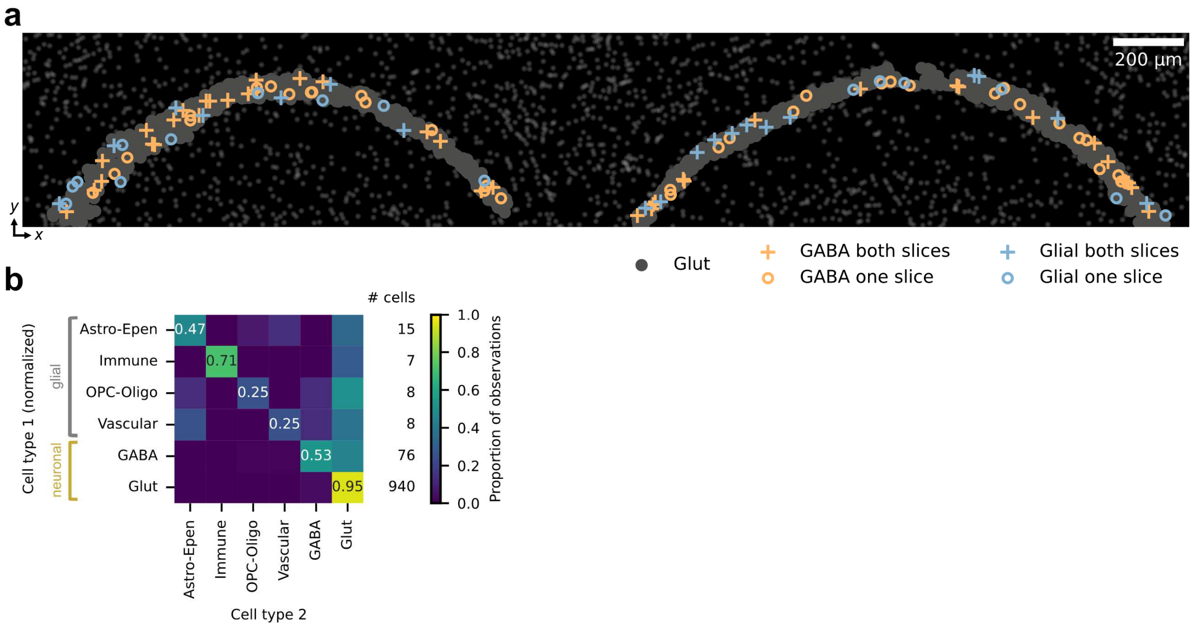

**Supplementary Figure 5: Cell overlaps in the CA1 region in the MERSCOPE mouse brain dataset detected through *in silico* vertical subslicing**. **a)** Cell segment map showing location of GABAergic and glial cell segments in CA1, marked as occurring consistently across both top and bottom virtual subslices or on one subslice only. **b)** Combinations of observed cell type assignments across top and bottom virtual subslices in CA1 cell segments (row-normalized), with many cell segments forming vertical doublets with the abundant glutamatergic neurons.

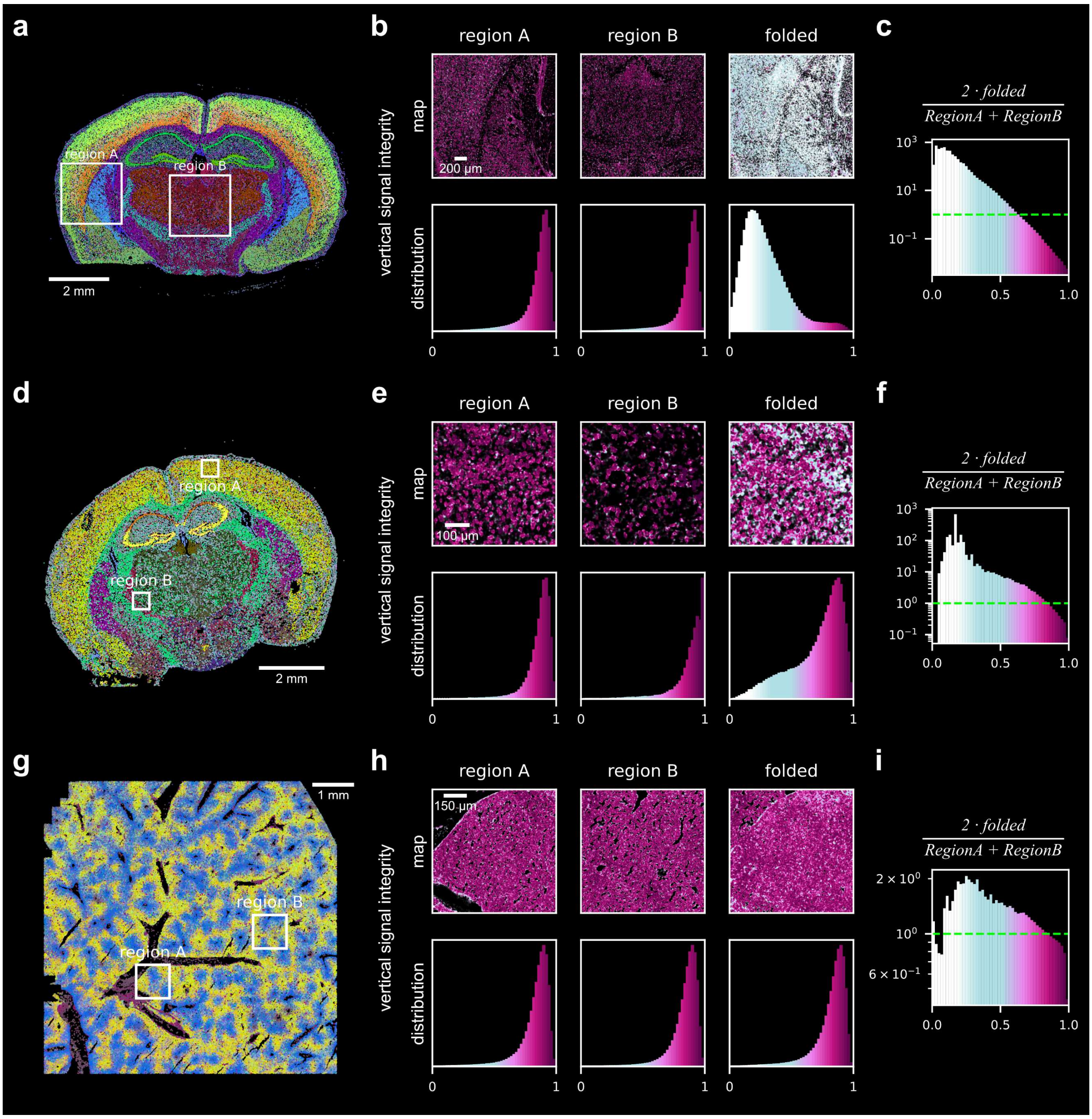

**Supplementary Figure 6: *In silico* synthetic tissue overlaps validate overlap detection model**. **a)** Transcript map of the Xenium mouse brain dataset colored using *ovrlpy* RGB embeddings, with regions highlighted for a synthetic tissue overlap (region A and B). **b)** Spatial map visualizations (top) and distributions (bottom) of vertical signal integrity for the regions A and B and the *in silico* synthetic tissue overlap of regions A and B. **c)** Log-ratio of distribution of vertical signal integrity in overlapped vs non-overlapped regions. Horizontal dashed line marks a ratio of 1:1. Same analysis for **d-f)** MERSCOPE mouse brain data set and **g-i)** MERSCOPE liver data set.

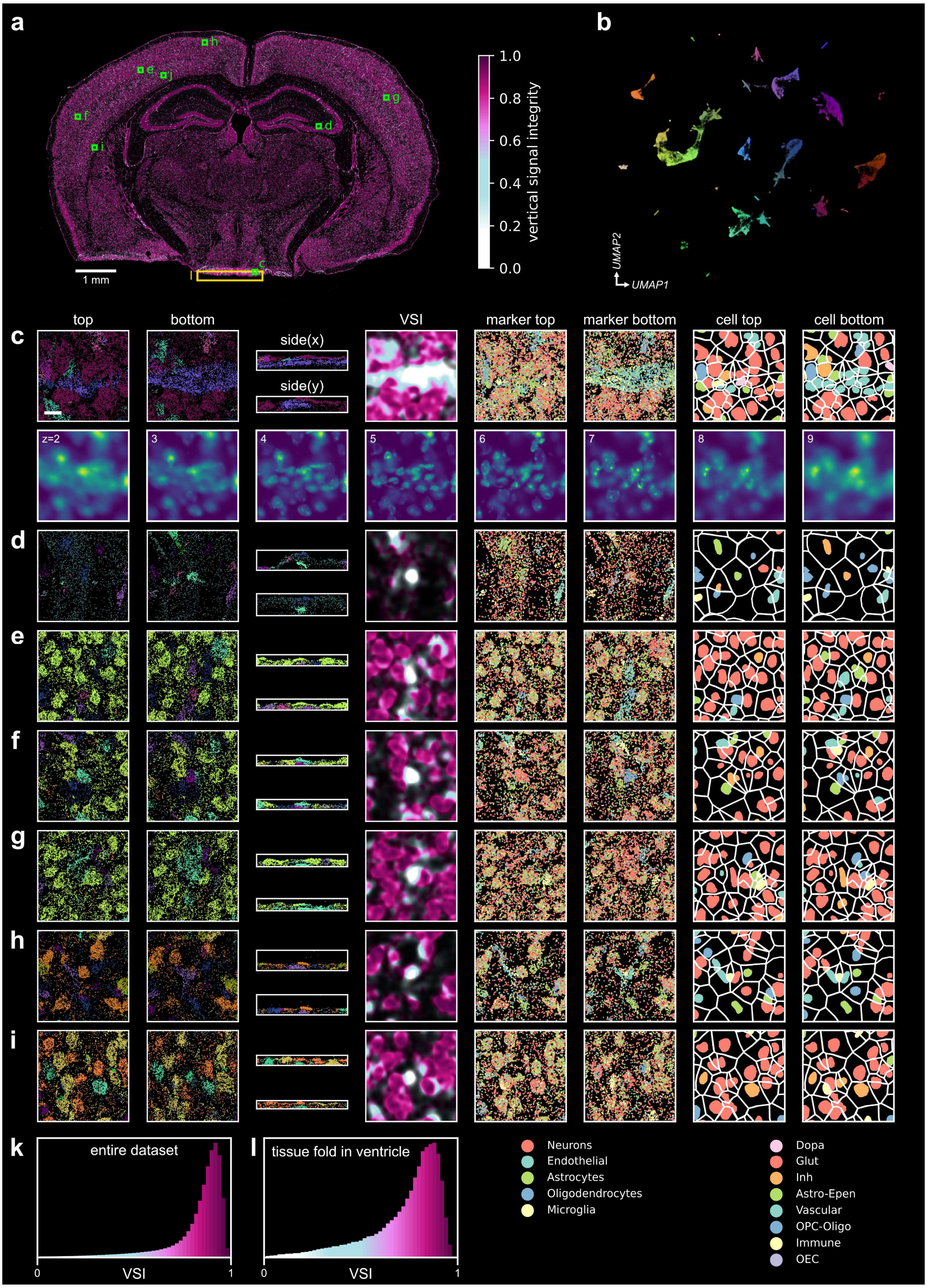

**Supplementary Figure 7: *Ovrlpy*-identified vertical doublets are consistent with supervised segmentation-based doublets identified in the Xenium mouse brain. a)** Vertical signal integrity (VSI) map of the Xenium mouse brain dataset, indicating 7 of the top vertical doublets identified by *ovrlpy*. **b)** UMAP and RGB embedding of the gene expression sampled at local maxima. Different cell types appear as separate clusters in the UMAP as well as the RGB embedding. **c-j)** Close-up of highlighted regions in panel a. The color-embedded transcripts highlight the different cell types and structures involved in spatial vertical doublets (column 1-3, RGB color-embedded as shown in panel b). VSI map of the highlighted region (column 4). Marker transcripts are based on the Xenium gene panel design (column 5, 6). Cell types are based on MapMyCells annotation of the top and bottom virtual subslice of nuclear transcripts (column 7, 8). Scale bar 15 µm. Distribution of VSI in the **k)** entire dataset and **l)** the tissue fold in the ventral region (highlighted in panel a).

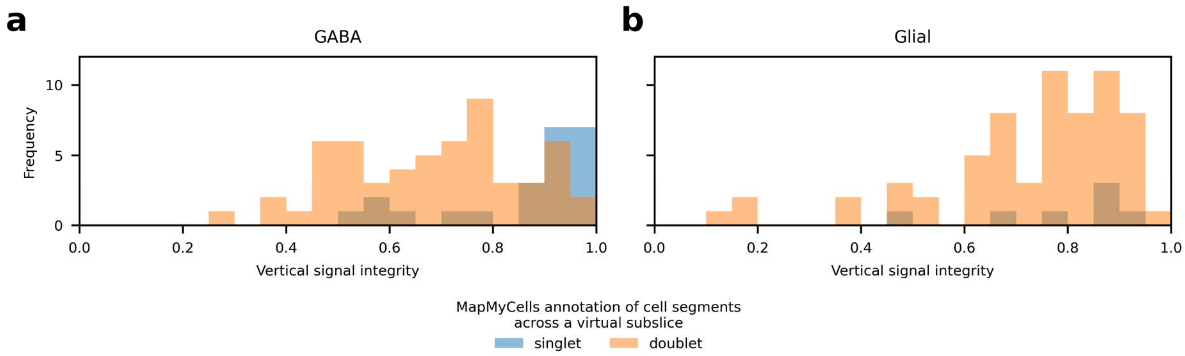

**Supplementary Figure 8: Vertical signal integrity distinguishes singlets and vertical doublets in mural CA1 cell segments**. Vertical signal integrity distribution in CA1, classified as vertical 'singlets' or 'doublets' based on a segmentation-based cell classification using the virtual subslicing strategy for **a)** GABAergic interneurons and **b)** glial cell segments. Singlets are underrepresented in both cell types and are more likely to exhibit a higher vertical signal integrity across virtual subslices in GABAergic interneurons (GABAergic cell segments: 23 singlets and 57 doublets with mean VSI of 0.86 and 0.68, $U = 1050$, $p < 0.001$; glial cell segments: 7 singlets and 65 doublets with mean VSI of 0.79 and 0.73, $U = 162$, $p = 0.22$; two-sided Mann-Whitney-U test).

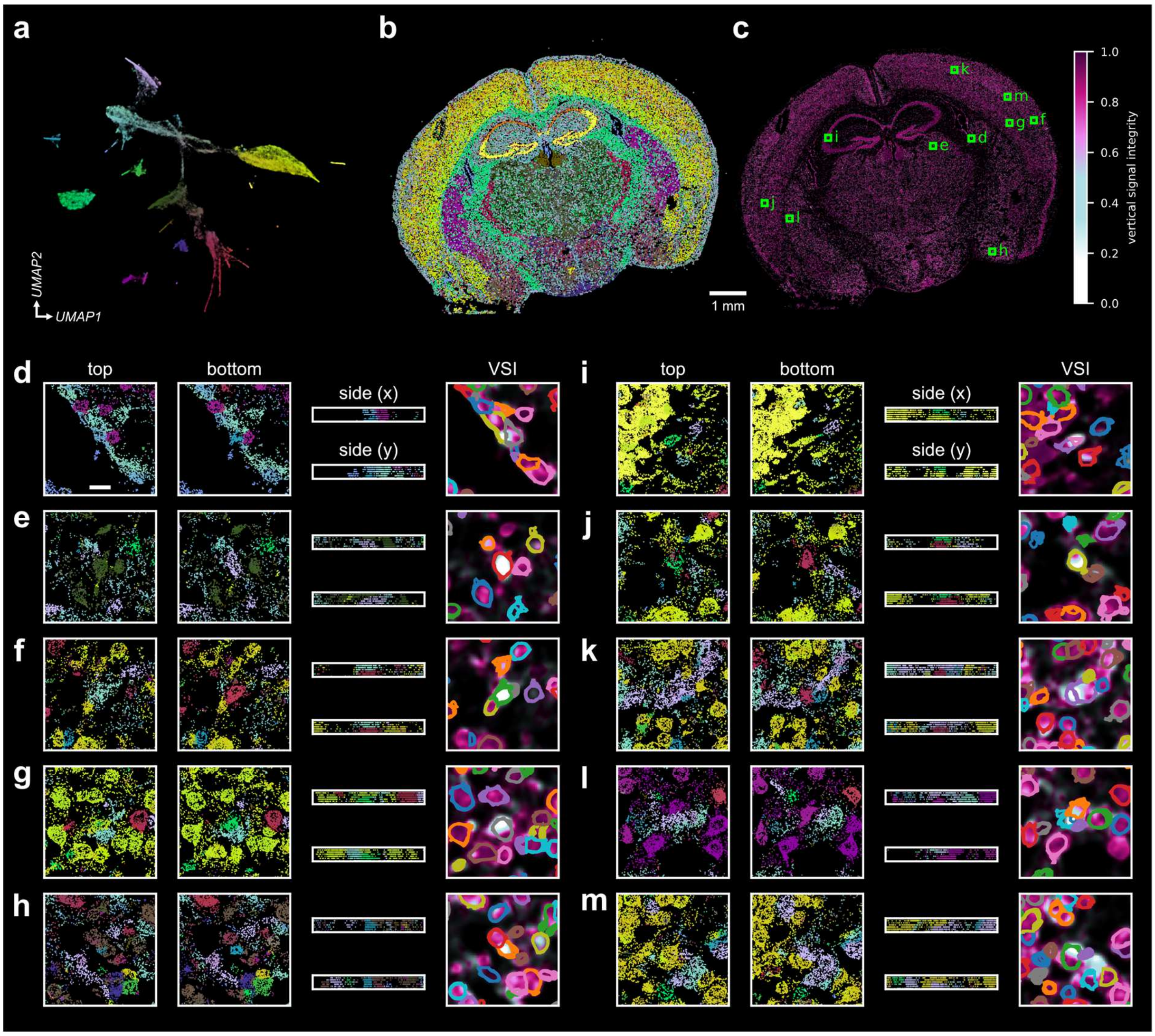

**Supplementary Figure 9: Vizgen's 3D segmentation fails to identify most cell overlaps detected by *ovrlpy* in the MERSCOPE mouse brain dataset**. **a)** UMAP and RGB embedding of the gene expression sampled at local maxima. **b)** Gene expression map of the MERSCOPE mouse brain dataset colored using *ovrlpy* RGB embeddings at local maxima sampling locations. **c)** Vertical signal integrity (VSI) map of the whole brain slice. The ten most prominent VSI local minima identified by *ovrlpy* are highlighted. **d-k)** Visualizations of vertical doublets. First three columns show transcript maps of top and bottom virtual subslices, and from the sides. Transcripts are colored by *ovrlpy* RGB color. Last column shows the VSI map in the corresponding region with overlayed segmentation masks for the different imaging layers. Scale bar 15 µm.

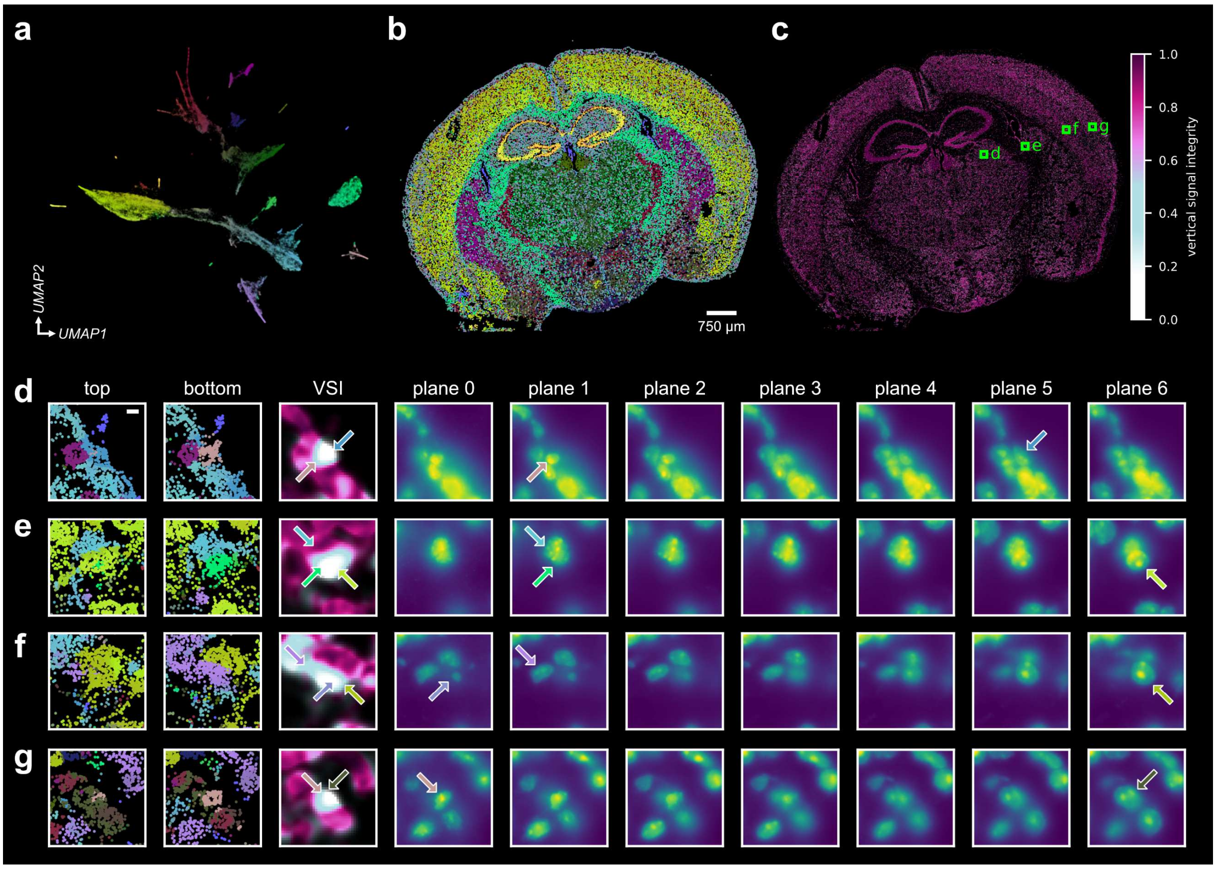

**Supplementary Figure 10: Nuclear stain supports *ovrlpy*-detected vertical doublets. a)** UMAP and RGB embedding of the gene expression sampled at local maxima. **b)** Gene expression map of the MERSCOPE mouse brain dataset colored using *ovrlpy* RGB embeddings at local maxima sampling locations. **c)** Vertical signal integrity (VSI) map of the whole brain slice. Four selected out of the ten most prominent VSI local minima identified by *ovrlpy* are highlighted and visualized in subsequent panels. **d-g)** Visualizations of vertical doublets. First 2 columns show transcript maps of top and bottom virtual subslices (colored by *ovrlpy* RGB embedding). Third column shows the VSI map in the corresponding region. The other columns correspond to the DAPI-stained images of multiple imaging focal planes (1.5 µm distance between adjacent planes). The arrows indicate nuclei participating in the overlap and are colored according to the RGB embedding in the first two columns. Scale bar 5 µm.

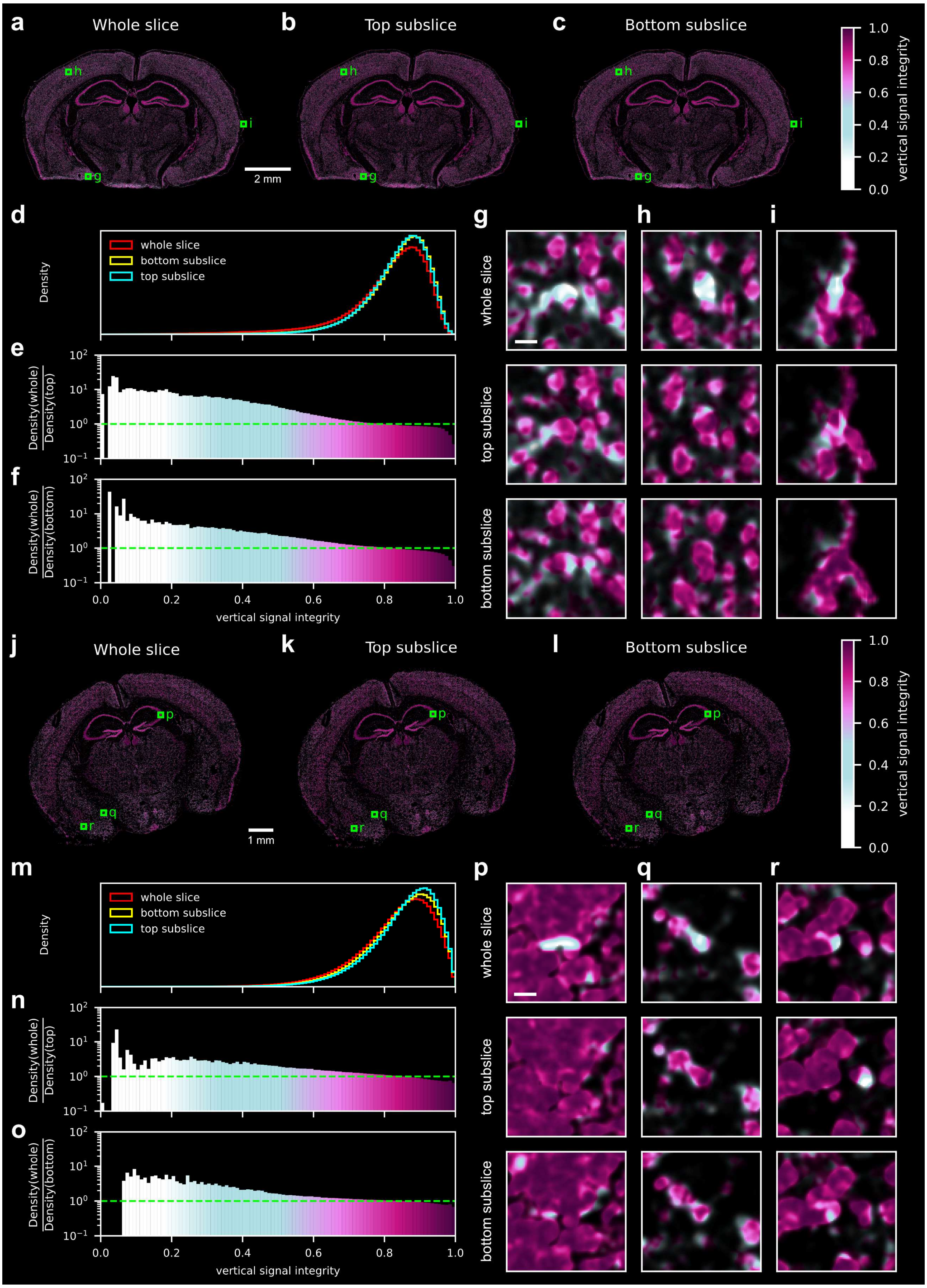

**Supplementary Figure 11: Subslicing a tissue section increases overall vertical signal integrity.** VSI map of **a)** the whole Xenium mouse brain, **b)** the top virtual subslice, and **c)** the bottom virtual subslice of the same sample. All sections have been downsampled to contain the same number of transcripts. **d)** The overall VSI of the bottom and top subslices are higher than in the whole tissue slice. **e)** Ratio of the VSI density between whole slice and top subslice and **f)** whole slice and bottom subslice demonstrate that low VSI scores are reduced in the virtual subsections. **g-i)** VSI of the top three vertical doublets identified in the whole tissue slice. Most vertical doublets are removed in the bottom and top subslices. Scale bar 15 µm. **j-r)** Same analysis as panel a-i repeated for the Vizgen MERSCOPE mouse brain sample.

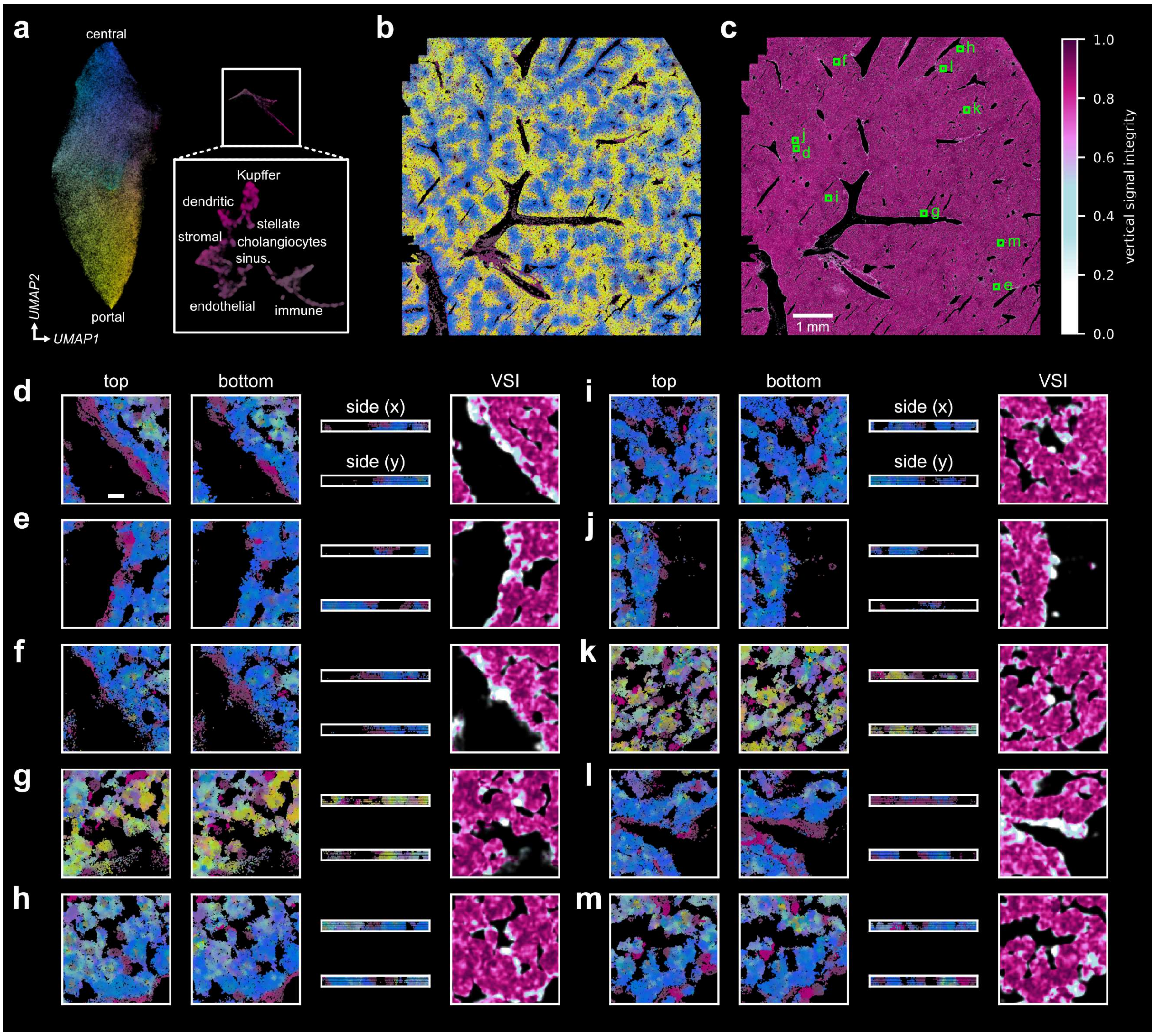

**Supplementary Figure 12: Overlaps identified in the MERSCOPE mouse liver dataset implicate many different cell types**. **a)** UMAP and RGB embedding of the gene expression sampled at local maxima. Different cell types appear as clusters in the UMAP and RGB embedding. Inset shows UMAP embedding based only on non-hepatocyte local maxima using the same colors. **b)** Gene expression map of the MERSCOPE liver dataset colored using *ovrlpy* RGB embeddings. **c)** Vertical signal integrity (VSI) map of the sample. Highlighted are the top ten vertical doublets identified by *ovrlpy*. **d-k)** Visualizations of vertical doublets; first three columns show transcript maps of top and bottom virtual subslices (colored according to panel a), last column shows the VSI map. Scale bar 15 µm.

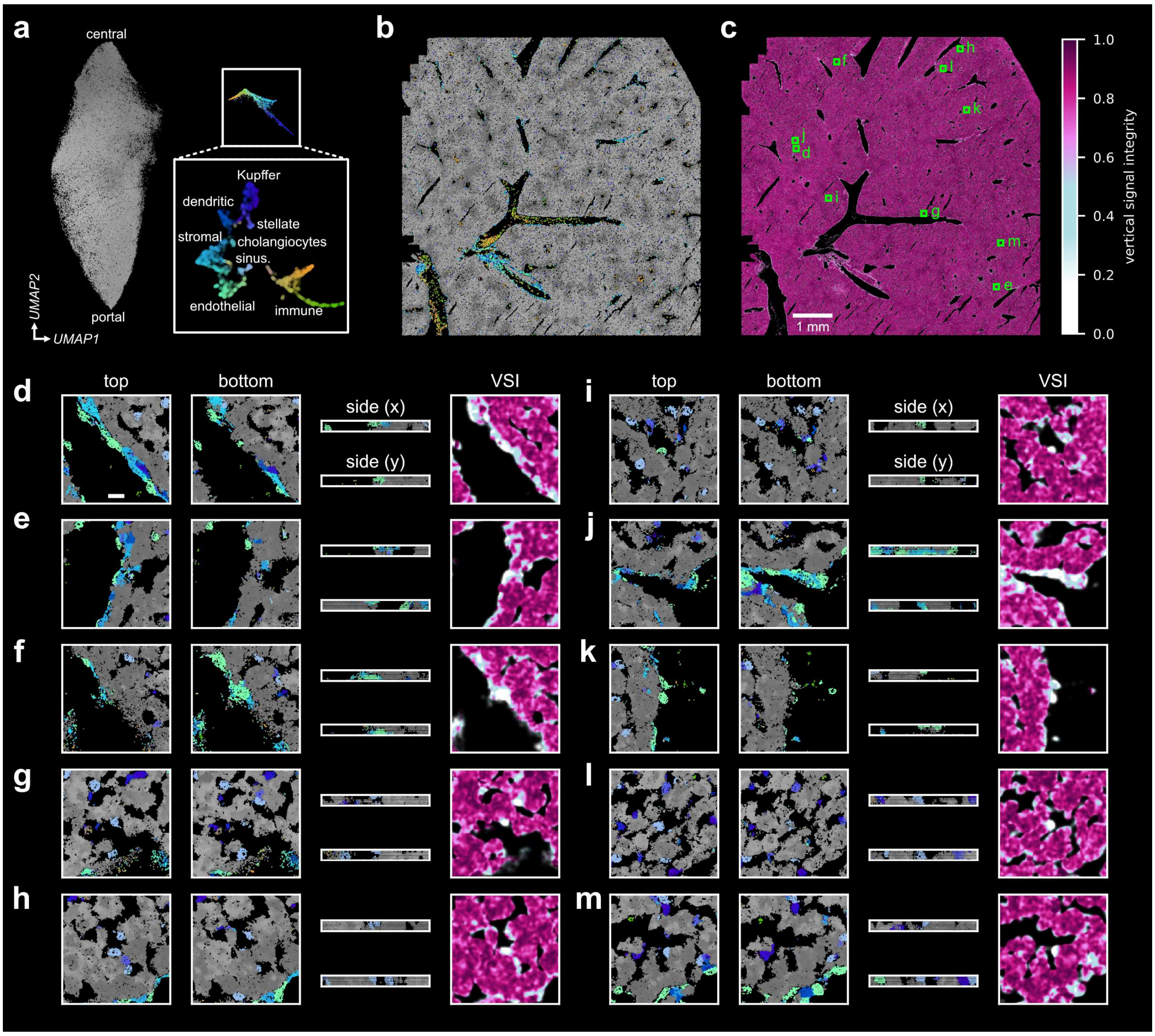

**Supplementary Figure 13: Overlaps identified in the MERSCOPE mouse liver dataset implicate many different cell types, using modified coloring for the gene expression embeddings to highlight the non-parenchymal cells**. **a)** UMAP and RGB embedding of the gene expression sampled at local maxima, using an added RGB model to colorize non-hepatocytes. Different cell types appear as clusters in the UMAP and RGB embedding. Inset shows UMAP and RGB embedding based only on non- parenchymal local maxima using the same colors. **b)** Gene expression map of the MERSCOPE liver dataset colored using *ovrlpy*'s RGB embedding (panel a). **c)** Vertical signal integrity (VSI) map of the sample. Highlighted are the top ten vertical doublets identified by *ovrlpy*. **d-k)** Visualizations of vertical doublets; first three columns show transcript maps of top and bottom virtual subslices (colored according to panel a), last column shows the VSI map. Scale bar 15 µm.

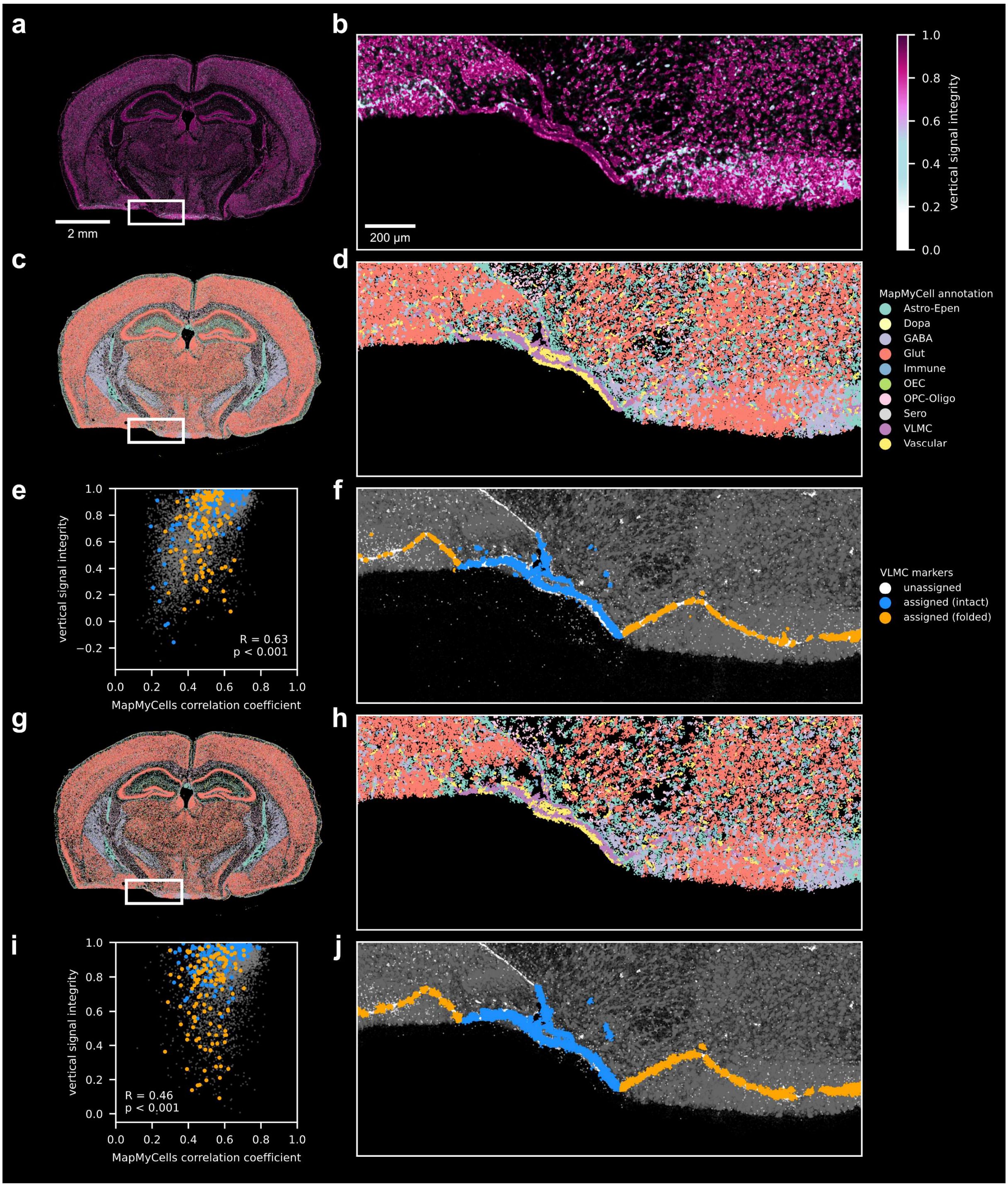

**Supplementary Figure 14: 3D segmentation followed by MapMyCells annotation fails to recapitulate the VLMCs in the tissue fold artifact**. VSI map of the **a)** whole tissue slice and **b)** highlighted ventral brain region containing the tissue fold artifact. **c)** Cell-type map of 3D Baysor segmentation cell centroids colored by the cell type family determined by MapMyCells. **d)** Transcript molecules are colored according to the cell type of their assigned cell segment. **e)** Vertical signal integrity (VSI) score, determined at cell centroids, correlates with MapMyCells' subclass correlation coefficient. All cells of the highlighted area in grey, VLMC in orange (within tissue fold) or blue (within intact artifact-free region). **f)** Transcript molecules in the region containing a folding artifact alongside an intact tissue region. VLMC markers: *Aldh1a2, Col1a1, Fmod, Slc13a4* **g-j)** Same analysis as c-f using Proseg for segmentation. Association of unfolded VLMCs with higher VSI compared to folded VLMCs was tested using a one-sided Mann-Whitney U test; Baysor: 130 folded VLMCs, 194 unfolded VLMCs, U = 19735, $p < 0.001$, Proseg: 129 folded VLMCs, 189 unfolded VLMCs, U = 20163, $p < 0.001$.

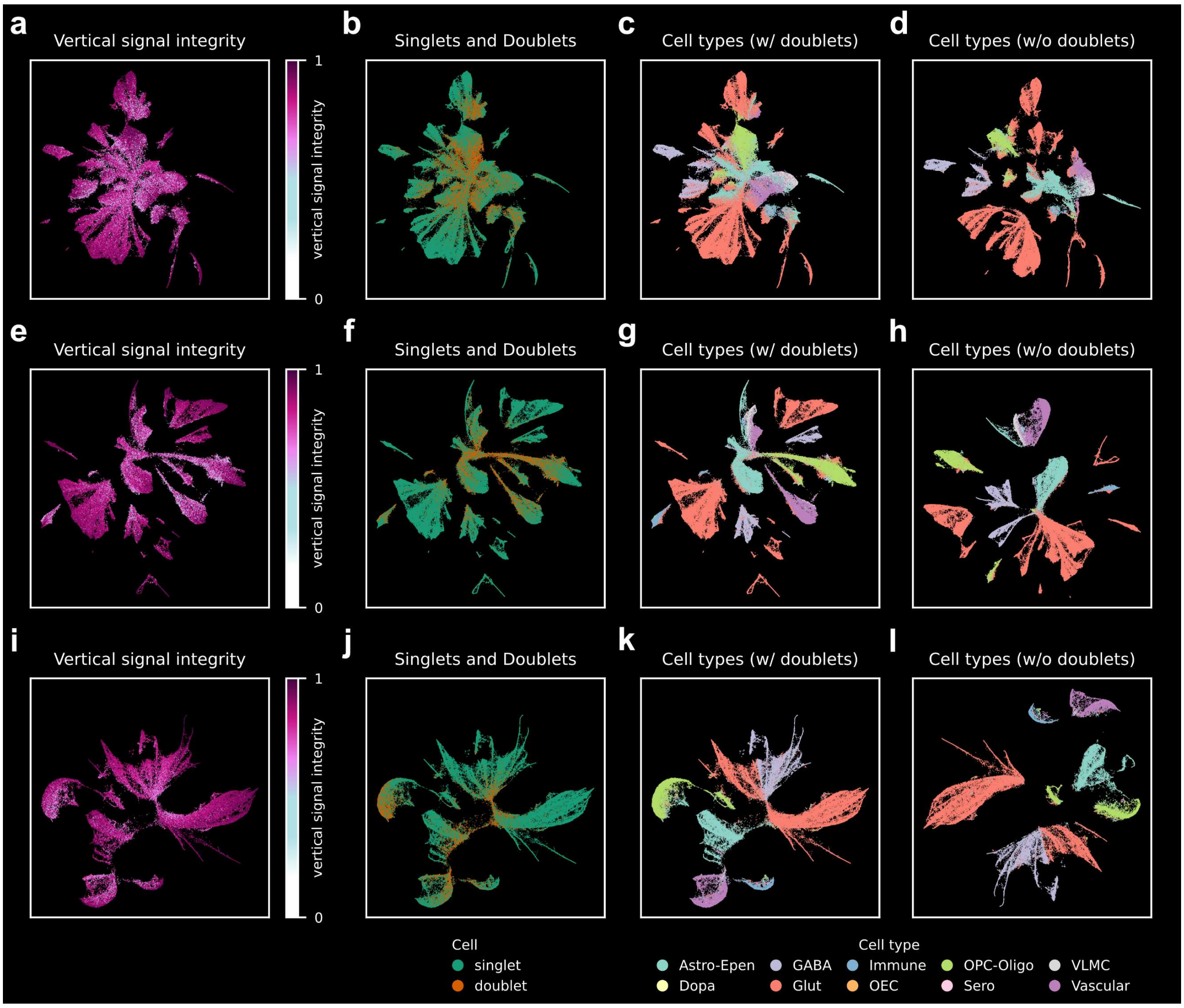

**Supplementary Figure 15: *Ovrlpy's* unsupervised vertical doublet filtering enhances cell-type separation in gene expression space. a-d)** UMAP embeddings of gene expression data from Xenium mouse brain cell segments colored by **a)** mean vertical signal integrity (VSI), **b)** singlets and vertical doublets (VSI threshold of 0.7; 162,033 cell segments of which 36,846 are vertical doublets), **c)** MapMyCells cell type annotations, **d)** MapMyCells cell type annotations (UMAP embedding after excluding vertical doublets). Same as a-d for **e-h)** the Xenium mouse brain with only nuclear transcripts (162,018 nuclei of which 36,831 are vertical doublets), and **i-l)** the nuclear segmented MERSCOPE mouse brain (83,505 nuclei of which 15,780 are vertical doublets).

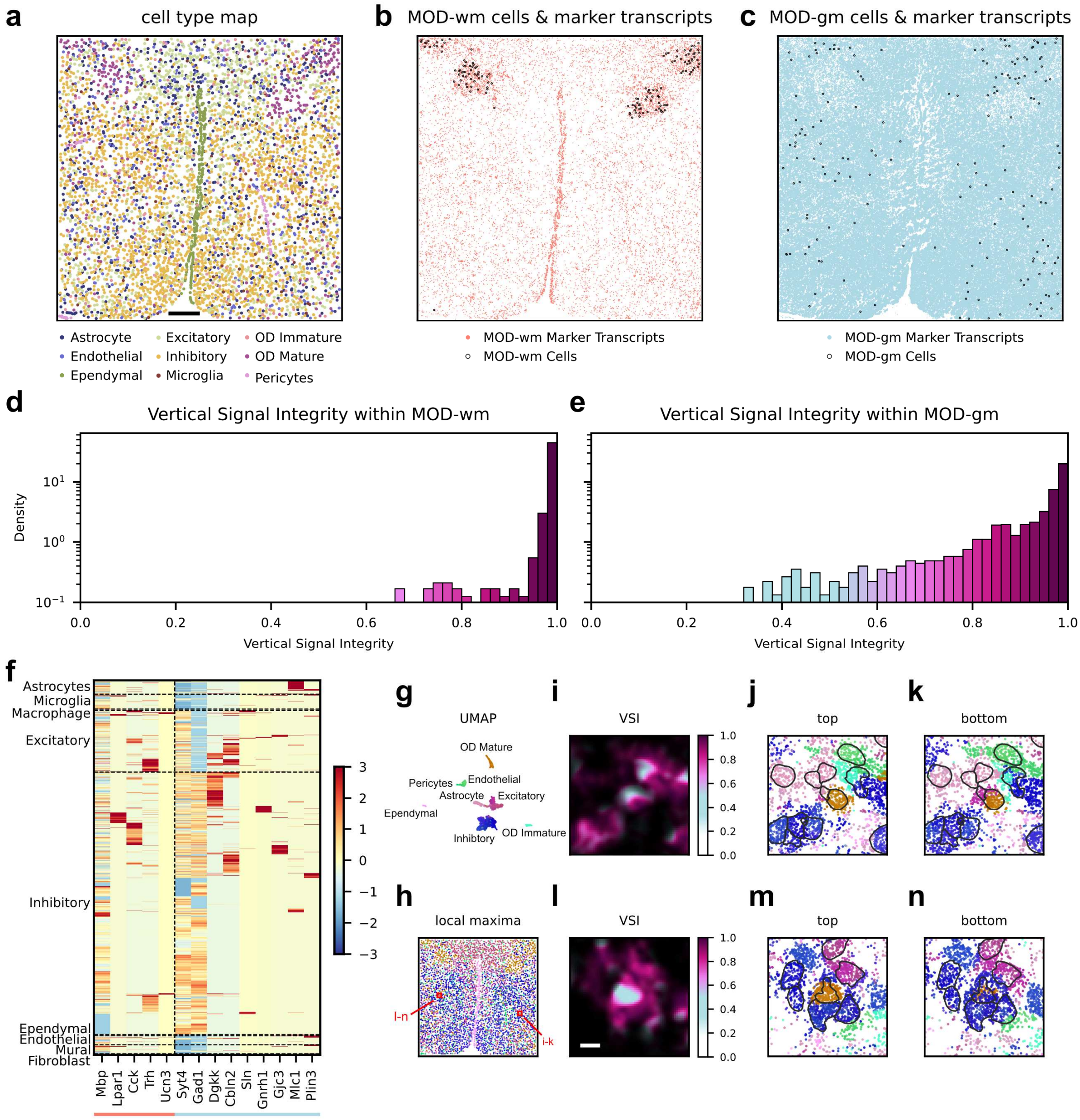

**Supplementary Figure 16: *Ovrlpy* identifies contaminations in spatial cell typing. a)** Cell types for the mouse hypothalamus dataset according to the original publication by Moffitt *et al.* (2018). Scale bar 200 µm. **b)** Location of mature oligodendrocyte (MOD) subtypes and their corresponding marker gene transcripts for MOD-wm, and **c)** MOD-gm as described by Singhal *et al*. (2024). Only cell segments not annotated as 'Ambiguous' (doublets according to Moffitt *et al.*) are shown. Notably, the proportion of these doublets for MOD-gm is larger than for MOD-wm (45% and 10% respectively, compared to 13% for all cell segments) implicating that the MOD-gm cell segments are potential subject to gene expression contamination from adjacent cell segments. VSI within **d)** MOD-wm and **e)** MOD-gm cell segments (excluding 'Ambiguous') demonstrates reduced VSI for MOD-gm cell segments. **f)** Expression of MOD-wm and MOD-gm marker genes in other cell types of the same tissue (matched scRNA-seq dataset) demonstrates systematic expression of MOD-gm markers in most neurons (*Syt4*), inhibitory neurons (*Gad1*), and astrocytes (*Mlc1*). **g)** *Ovrlpy* UMAP embedding and **h)** corresponding local maxima locations. Examples of annotated MOD-gm cell segments overlapping with **i-k)** excitatory and **l-n)** inhibitory neurons. Scale bar 10 µm. gm: gray matter, wm: white matter

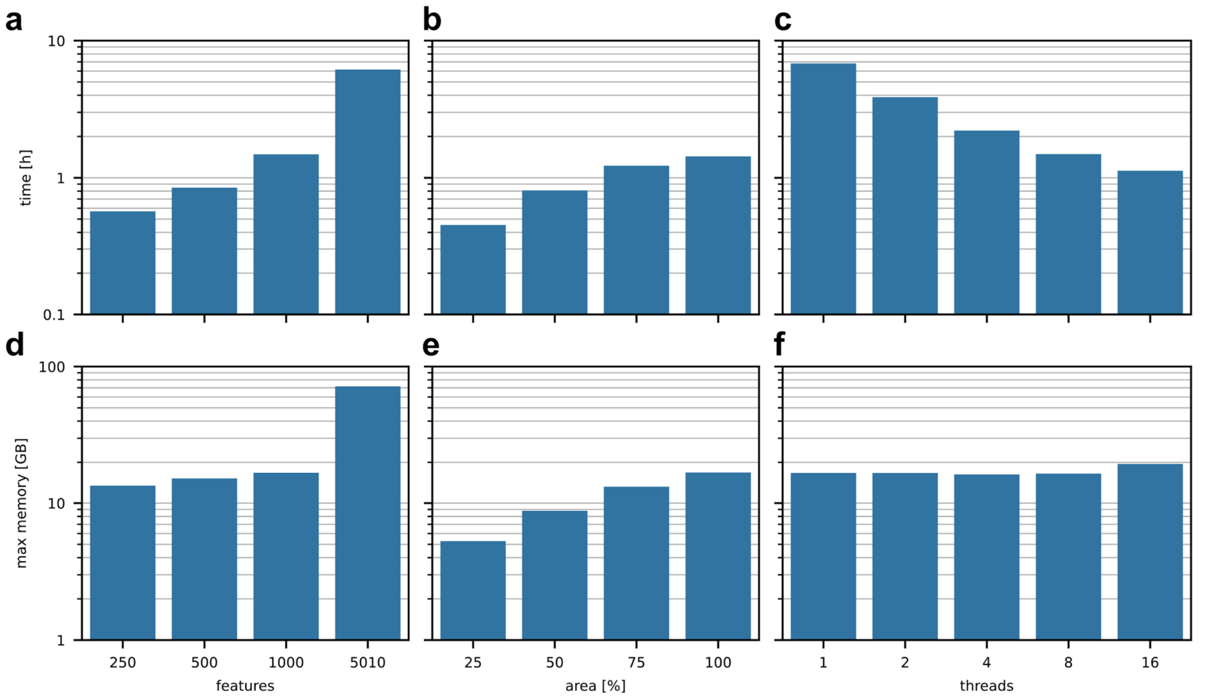

**Supplementary Figure 17: Runtime and memory usage of *ovrlpy*. a-c)** Runtime and **e-f)** max. memory usage when processing the Xenium Prime 5k mouse pup downsampled to 1,000 genes (unless otherwise specified) resulting in 125,756,304 transcripts and a sample area of 272.6 $mm^2$ using 8 threads by default for **a, d)** varying number of genes, **b, e)** subset of the area, and **c, f)** processing with varying number of threads.
